# Critical illness expands a transcriptionally distinct hypometabolic CD8^+^ T effector program associated with respiratory failure and mortality

**DOI:** 10.64898/2026.05.28.728555

**Authors:** Casey M. Nichols, Erin L. Mwizerwa, Chooi Ying Sim, Sarah N. Obeidalla, Jacqueline-Yvonne Cephus, Caroline E. Roe, Jonathon M. Irish, Dawn C. Newcomb, V. Eric Kerchberger, Julie A. Bastarache, Jeffrey C. Rathmell, Lorraine B. Ware, Matthew T. Stier

## Abstract

Immune dysfunction is a major driver of morbidity and mortality in critical illness syndromes including sepsis. Specifically, CD8^+^ T cell dysfunction has been linked to organ failure and death. To characterize the immune substructure of circulating CD8^+^ T cells in critical illness at high dimension, we used single-cell RNA sequencing of peripheral blood CD8^+^ T cells from 38 critically ill patients and 9 healthy controls. We annotated seven CD8^+^ T cell clusters, which included a CD8^+^ effector subset, termed T effector state 2 (T_Eff-2_), that was only present in critically ill patients and associated with more severe respiratory failure and higher mortality. T_Eff-2_ showed effector activation and inflammatory stress conditioning yet had markedly reduced metabolic transcripts without canonical features of exhaustion. Trajectory analyses positioned T_Eff-2_ as a terminal CD8^+^ T effector cell fate driven in part by *DDIT4* and *DUSP1*, which negatively regulate mTOR and MAPK signaling, respectively. Interestingly, this transcriptional program was indistinguishable by classical protein cytometry methods. These results, including the mortality association, were validated in a larger (n=91) independent external cohort of critically ill patients with sepsis. In summary, T_Eff-2_ represents a latent transcriptional program that delineates a clinically high-risk CD8^+^ T cell state in critical illness.

## Introduction

Critical illness conditions such as sepsis, acute respiratory distress syndrome (ARDS), and trauma impose a substantial public health burden, with an in-hospital mortality of 22% globally (1). In the United States, over 5 million patients require intensive care unit (ICU) admission annually (2), with costs exceeding US$108 billion (3). The burden of critical illness extends well beyond acute mortality; survivors suffer lasting physical and cognitive impairment and disability at rates exceeding matched hospitalized controls (4, 5). Despite advances in supportive care, ICU mortality remains high (6), and no targeted therapies have demonstrated consistent survival benefits for critical illness syndromes such as sepsis (7).

Immune dysfunction is a hallmark of critical illness and contributes substantially to morbidity and mortality (8–12). T lymphocytes are profoundly affected, with increased apoptosis, marked lymphopenia, and an exhausted-like state with checkpoint regulator upregulation and impaired cytokine production (13). These changes are linked to secondary infections, viral reactivation, persistent organ dysfunction, and death (12, 14–16). CD8^+^ T cell impairment, in particular, is associated with increased organ failure and mortality (12, 17–21). However, most studies have analyzed CD8^+^ T cells as a bulk population, leaving the clinical significance of specific CD8^+^ T cell subsets poorly understood in critical illness.

Biological heterogeneity is a major barrier to therapeutic progress in critical illness syndromes. Over 100 randomized clinical trials in sepsis have failed to demonstrate consistent benefit, at least in part because trial populations do not discern between biologically distinct patient subgroups (22). Indeed, bulk transcriptomic approaches have identified reproducible immune endotypes in sepsis with divergent outcomes and treatment responses, establishing that potentially clinically actionable biological substructure exists within these syndromes (23–26). Single-cell transcriptomics has further resolved disease-associated immune cell states with prognostic significance not detectable by bulk profiling (27–30). In sepsis, these high-dimensional approaches have identified an immunosuppressive monocyte state (MS1) (27) and linked emergency granulopoiesis-driven neutrophil expansion to a high-mortality endotype (28). The CD8^+^ T cell compartment has received comparatively less assessment, leaving its heterogeneity and clinical associations largely uncharacterized.

In this study, we used single-cell RNA sequencing (scRNA-seq) to profile circulating CD8^+^ T cells from critically ill patients with and without sepsis and healthy controls. We identified a novel CD8^+^ effector subset, T_Eff-2_, with a distinct hypometabolic phenotype and mixed activation and cell stress features. T_Eff-2_ frequency was associated with respiratory failure and mortality, and transcriptomic features of T_Eff-2_ were partially shared with bronchoalveolar lavage (BAL) tissue resident memory CD8^+^ T cells in pneumonia patients. Finally, T_Eff-2_ frequency and phenotype were validated in an independent external sepsis cohort, where T_Eff-2_ abundance associated with 90-day mortality.

## Results

### Critical illness expands a transcriptionally distinct CD8^+^ T cell effector population

To investigate the CD8^+^ T cell compartment in critical illness, we leveraged our previously reported 644,147 cell scRNA-seq peripheral blood mononuclear cells (PBMCs) database from 9 healthy controls (HC; 44% male, median age 47, IQR 34-58) recruited from the community and 38 critically ill patients (CI; 55% male, median age 65, IQR 55-73) recruited from the medical and surgical ICUs at Vanderbilt University Medical Center with blood collected 48-72 hours post-admission (Fig. 1A, Table 1) (30). The ICU cohort included 19 patients with sepsis and 19 patients with mixed critical illness not due to sepsis. We restricted analyses to CD8^+^ T cells excluding the low abundance non-traditional mucosal-associated invariant T (MAIT) cells, and then re-clustered and annotated using canonical transcriptional markers, identifying 7 distinct cell populations (Fig. 1B-D, Supplemental Fig. 1A). We identified major known circulating CD8 T cell subsets including naïve (T_N_), central memory (T_CM_), effector memory (T_EM_), KIR^+^ (T_KIR_), and CMC1^+^ (T_CMC1_) cells. Consistent with previously reported findings, we observed decreased T_N_ (31) and T_KIR_ (32) CD8^+^ T cells in CI compared to HC (Fig. 1E, F, Supplemental Fig. 1B). We also identified two effector subsets, which we termed T effector state 1 (T_Eff-1_) and T effector state 2 (T_Eff-2_). T_Eff-1_ were present in both HC and CI, whereas T_Eff-2_ were only observed in appreciable frequencies in CI patients (mean frequency 0.1% HC, 8.1% CI; Fig. 1E, Supplemental Fig. 1B). Frequencies of T_Eff-2_ in CI patients varied widely, from <0.1% to >50% of all CD8 T cells (Fig. 1F), and were similar between CI non-septic patients (CI-NS) and CI patients with bacterial sepsis (CI-Sep), indicating this effector state is a feature of critical illness broadly rather than sepsis specifically (Fig. 1G).

**Figure 1:**
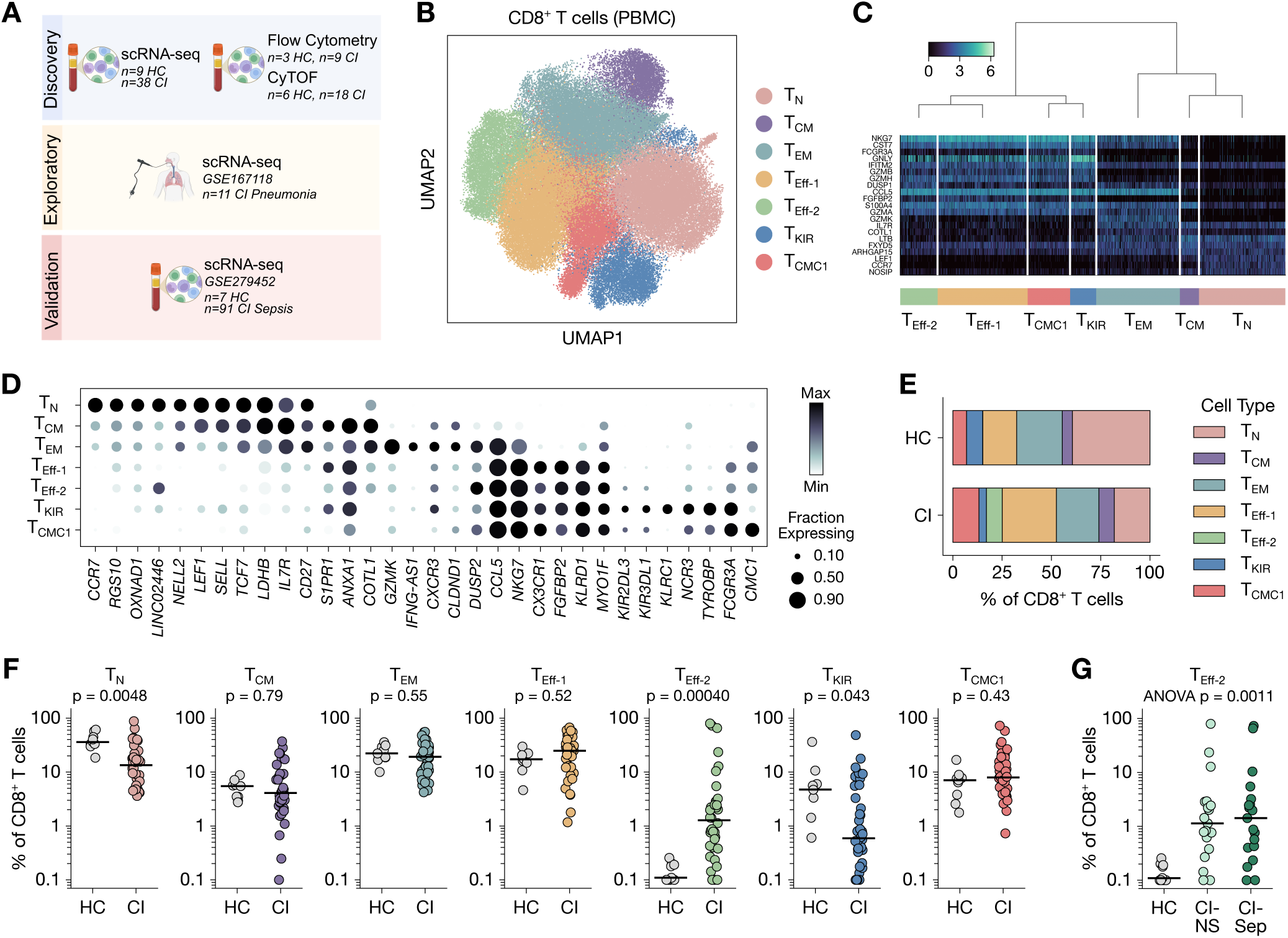
Heterogeneity of circulating CD8^+^ T cells in critical illness. **(A)** Cohort schematic. Created in BioRender. Nichols, C. (2026) https://BioRender.com/eilqhpx. **(B)** UMAP visualization of CD8^+^ T cells from PBMC of HC and CI patients annotated by cell cluster. **(C)** Top defining genes for each cell cluster in B. Cluster order sorted hierarchically by gene expression profile. **(D)** Normalized scaled gene expression of selected cluster-defining genes. **(E)** Proportion of each T cell cluster among all CD8^+^ T cells in HC and CI patients. **(F)** Per participant proportion of each T cell cluster among all CD8^+^ T cells in HC and CI patients. **(G)** Per participant proportion of T_Eff-2_ cluster between CI non-septic (CI-NS) and CI septic (CI-Sep) patients. Statistical analysis was performed using the Mann-Whitney U testing with Benjamini-Hochberg adjustment **(C)** and the Empirical Bayes moderated T-test **(F)** or ANOVA **(G)**. Each datapoint represents an individual research participant.

### T_Eff-2_ frequency is associated with severe hypoxemic respiratory failure

To understand the clinical significance of T_Eff-2_, we next analyzed associations between T_Eff-2_ frequency and severity of illness and organ dysfunction. T_Eff-2_ frequency was not associated with admission Acute Physiology and Chronic Health Evaluation II (APACHE II) score, suggesting it was not broadly linked to composite illness severity as measured at ICU admission (Fig. 2A). Among individual organ failures, high T_Eff-2_ frequency was associated with more severe hypoxemic respiratory failure as measured by PaO_2_/FiO_2_ ratio (alveolar-gas exchange; n=23 samples with available arterial blood gas data) (Fig. 2A). However, T_Eff-2_ frequency did not correlate with measures of cardiovascular (norepinephrine equivalent dose, NEE; lactate; blood pH), renal (creatinine), or neurologic (Glascow Coma Scale, GCS) injury, suggesting a respiratory-centric relationship (Fig. 2A). Consistent with this, median T_Eff-2_ frequency increased stepwise from no hypoxemic respiratory failure (PaO_2_/FiO_2_ >300) to mild (201–300), moderate (101–200), and severe (<100) hypoxemic respiratory failure (Fig. 2B). This relationship with hypoxemic respiratory failure was unique to T_Eff-2_ (Fig. 2C, D).

**Figure 2:**
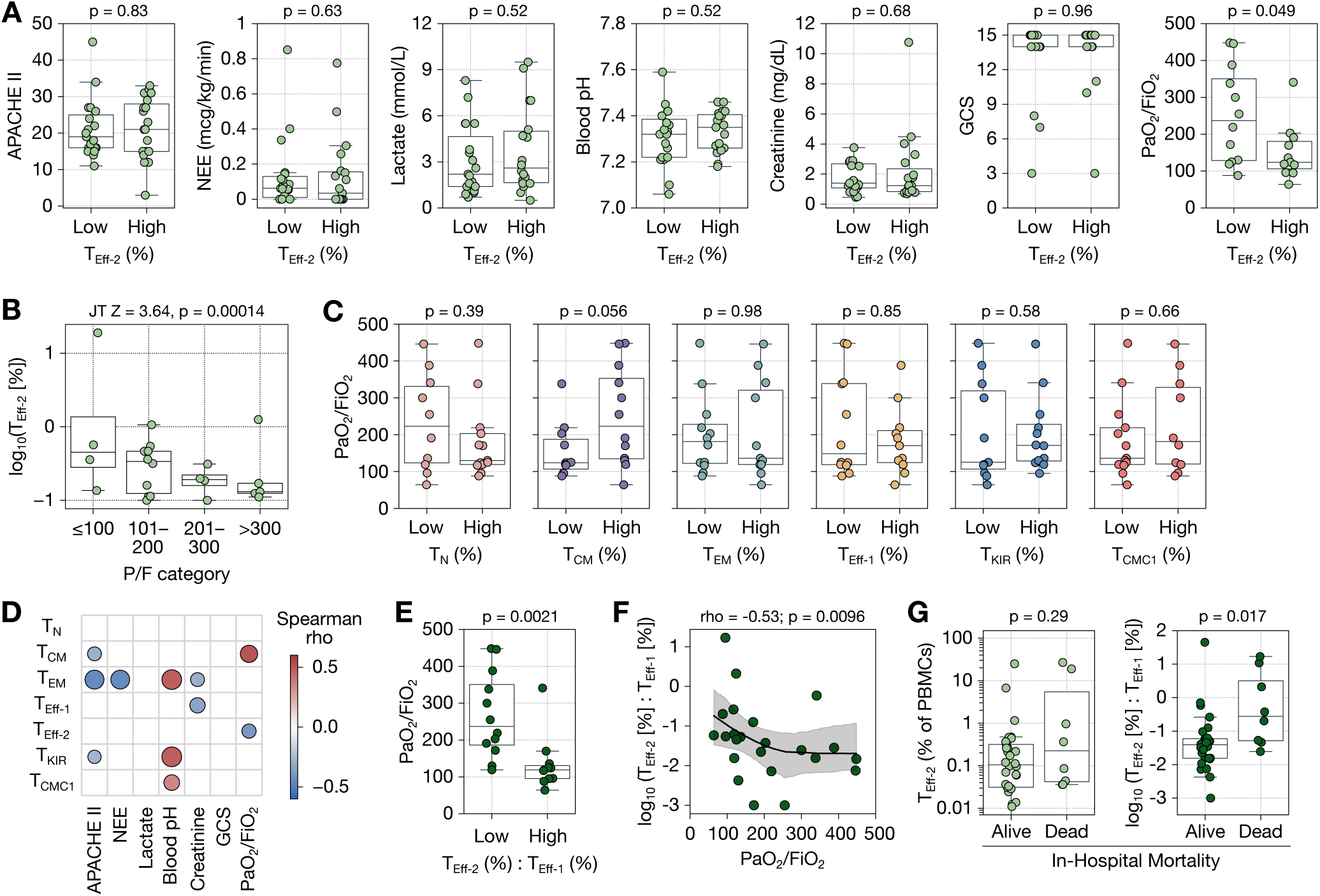
CD8^+^ T_Eff-2_ frequency is associated with respiratory failure. **(A)** ICU clinical variables within the first 24 hours of admission dichotomized by CD8^+^ T_Eff-2_ proportion of all PBMC above (High) or below (Low) the median proportion. **(B)** Log-normalized proportion of CD8^+^ T_Eff-2_ among PBMC stratified by severity of respiratory impairment measured by PaO_2_/FiO_2_ ratio. **(C)** PaO_2_/FiO_2_ ratio dichotomized by CD8^+^ T cell cluster proportion among all PBMC above (High) or below (Low) the median proportion. **(D)** Spearman rank correlation of CD8^+^ T cell cluster proportion and clinical variables. Associations only shown for p < 0.10. **(E)** PaO_2_/FiO_2_ ratio dichotomized by the ratio of T_Eff-2_:T_Eff-1_ above (High) or below (Low) the median proportion. **(F)** Spearman rank correlation showing the association between T_Eff-2_:T_Eff-1_ ratio and PaO_2_/FiO_2_ ratio. A locally weighted scatter-plot smoothing (LOWESS) curve was fitted and bootstrapped 95% confidence intervals are shaded. **(G)** T_Eff-2_ proportion of all PBMC and T_Eff-2_:T_Eff-1_ ratio dichotomized by in-hospital mortality status. Statistical analysis was performed using the Mann-Whitney U test **(A,C,E,G)**; the Jonckheere-Terpstra test **(B)**; and the spearman rank correlation test **(E-F)**. Boxplots show the median with boxes bounded by interquartile range (IQR; 25–75th percentile) and whiskers extending to the minimum and maximum values within 1.5×IQR. Each datapoint represents an individual research participant.

Because T_Eff-1_ and T_Eff-2_ represent distinct effector states, we reasoned that the balance between them may capture the skewing of the effector CD8^+^ T cell compartment towards T_Eff-2_ better than T_Eff-2_ frequency alone. Similar to T_Eff-2_ frequency alone, an increased T_Eff-2_:T_Eff-1_ ratio was associated with more severe respiratory impairment (Fig. 2E, F). Additionally, an elevated T_Eff-2:_T_Eff-1_ ratio was significantly associated with in-hospital mortality (Fig. 2G), while higher T_Eff-2_ frequency alone had numerically greater mortality that did not reach statistical significance (Fig. 2G). Collectively, these data identify CD8^+^ T_Eff-2_ as a critical illness-expanded subset linked selectively to hypoxemic respiratory failure and in-hospital mortality.

### T_Eff-2_ is defined by a hypometabolic yet activated and stressed-conditioned transcriptional program

We next sought to characterize the T_Eff-2_ transcriptional program. We performed univariate linear modeling (ULM) of Hallmark pathways across CD8^+^ effector lineage cells including T_EM_, T_Eff-1_, T_Eff-2_, T_KIR_, and T_CMC1_, (Fig. 3A, Supplemental Table 1). T_Eff-2_ demonstrated broad downregulation of most Hallmark pathways relative to other lineages. Metabolic pathways were universally decreased in T_Eff-2_ compared to other effector lineages, including glycolysis, oxidative phosphorylation, fatty acid metabolism, xenobiotic metabolism, mTOR/mTORC1 signaling, and MYC targets. Among the few positively enriched pathways in T_Eff-2_ were those associated with inflammatory signaling, including TNFA_SIGNALING_VIA_NFKB.

**Figure 3:**
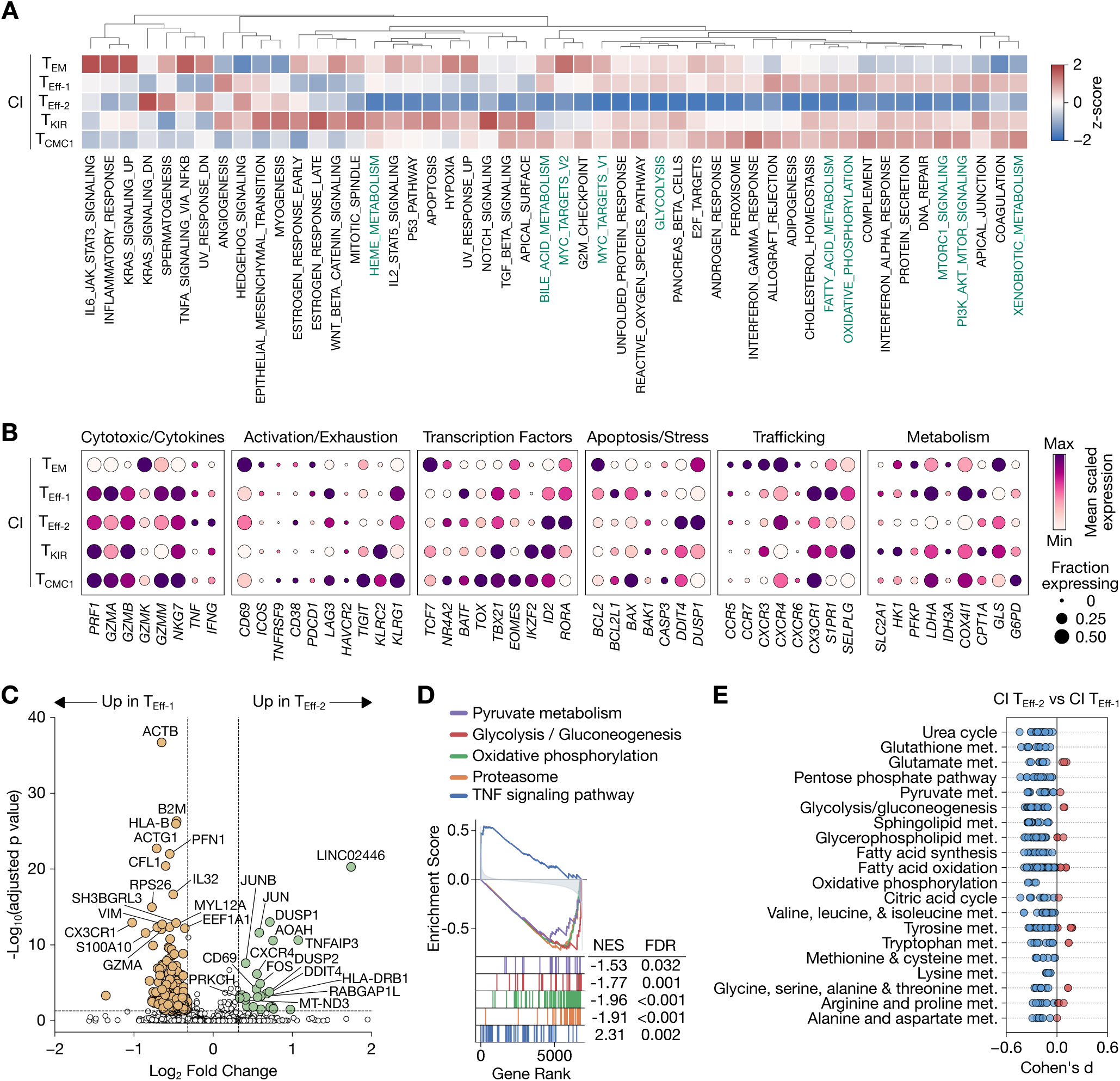
CD8^+^ T_Eff-2_ are hypometabolic with a mixed activated and stressed phenotype. **(A)** Univariate linear modeling (ULM) on a per donor basis of Hallmark pathways in CI CD8^+^ T cell subsets. Metabolism-associated pathways are highlighted in green. **(B)** Normalized scaled gene expression of selected functional and regulatory genes in CI CD8^+^ T cell subsets. **(C)** Differential gene expression analysis of CI T_Eff-2_ compared to CI T_Eff-1_. **(D)** Gene set enrichment analysis (GSEA) of selected KEGG pathways comparing CI T_Eff-2_ to CI T_Eff-1_. **(E)** Flux balance analysis via Compass of major metabolic reaction pathways. Each data point in **(E)** represents an individual metabolic reaction within the pathway and the effect sizes are represented by Cohen’s d. Statistical analysis was performed using the Mann-Whitney U with Benjamini-Hochberg adjustment **(C, E)** and the permutation-based Kolmogorov–Smirnov statistic with Benjamini–Hochberg correction **(D)**. NES, normalized enrichment score; FDR, false discovery rate.

To explore gene-level resolution of the T_Eff-2_ state, we examined normalized scaled expression of individual genes in T_Eff-2_ relative to other effector lineage populations (Fig. 3B). Consistent with an effector phenotype, T_Eff-2_ expressed cytotoxic mediators (*PRF1*, *GZMA*, *GZMB, GZMM*) and inflammatory cytokines (*TNF*, *IFNG*) but was skewed toward inflammatory cytokines over cytotoxic mediators compared to T_Eff-1_. T_Eff-2_ showed low expression of exhaustion-associated transcripts (*PDCD1*, *HAVCR2*, *TIGIT*, *TOX*). Among transcription factors, T_Eff-2_ highly expressed *ID2*, which promotes effector fates, and *RORA*. Moreover, T_Eff-2_ showed substantial evidence of inflammatory and metabolic stress, including increased expression of *DDIT4*, which suppresses the master metabolic regulator mTORC1, and *DUSP1,* which negatively regulates MAPK signaling. Consistent with the broad metabolic suppression identified by pathway analysis, T_Eff-2_ had reduced expression of numerous metabolic genes, including glycolytic enzymes (*HK1*, *PFKP*, *LDHA)* and mitochondrial TCA and respiratory complex components (*IDH3A*, *COX4I1)*. Differential gene expression analysis comparing T_Eff-2_ directly to T_Eff-1_ further corroborated a mixed activated and stressed phenotype (Fig. 3C, Supplemental Table 2). T_Eff-2_ had increased expression of the early activation marker *CD69*, the AP-1 transcription factor *JUN*, the TNF-induced NF-κB negative regulator *TNFAIP3*, *DUSP1,* and *DUSP2*. Among the most downregulated genes in T_Eff-2_ versus T_Eff-1_ were the cytotoxic mediator *GZMA* and *CX3CR1*. Together, these transcriptional features suggest T_Eff-2_ is a non-exhausted effector state conditioned by inflammatory stress and characterized by broad metabolic suppression.

To further compare T_Eff-2_ and T_Eff-1_, we performed gene set enrichment analysis using KEGG library pathways (Fig. 3D and Supplemental Table 3). TNF signaling was significantly enriched in T_Eff-2_, consistent with inflammatory cytokine exposure. Metabolic pathways including glycolysis, oxidative phosphorylation, and pyruvate metabolism were significantly depleted, supporting a transcriptionally hypometabolic state. Proteasome transcripts were also depleted in T_Eff-2_, suggesting broader suppression of biosynthetic and protein turnover programs. To gain reaction-level metabolic flux resolution of the T_Eff-2_ metabolic program, we applied *in silico* flux balance analysis (33), which models the stoichiometry of the Recon2 human metabolic network to estimate metabolic flux (Fig. 3E, Supplemental Fig. 2, Supplemental Table 4). T_Eff-2_ showed reduced predicted metabolic flux across all pathways and nearly all metabolic reactions relative to T_Eff-1_, providing convergent evidence of a profoundly hypometabolic state.

### Gene regulatory networks support both activation and metabolic restraint in T_Eff-2_

To investigate transcription factor programs underlying the mixed activated and hypometabolic identity of T_Eff-2_, we performed ULM of the CollecTRI transcription factor network, revealing distinct gene regulatory activity across effector-lineage CD8^+^ T cell subsets (Fig. 4A, Supplemental Table 5). Compared with T_Eff-1_, T_Eff-2_ showed increased activity of activation-associated transcription factors *NFATC3* and *REL*. Additionally, we observed increased activity of *FOXO3* and TGF-B effectors (*SMAD3*, *KLF10*), further supporting a hypometabolic state (Fig. 4B). To identify genes under coordinated control by T_Eff-2_-associated transcriptional regulators, we integrated regulator-target relationships with the T_Eff-2_ versus T_Eff-1_ delta-ULM-derived effect size. The resulting T_Eff-2_ regulatory network showed convergent net activation of AP-1 transcription factors *FOS* and *JUN*, inflammatory cytokines *TNF* and *IFNG*, the metabolic deacetylase *SIRT1*, transcriptional co-activator *CREBBP*, and the senescence-associated cell cycle inhibitor *CDKN2A* (Fig. 4C). These data collectively provide evidence of a coordinated transcriptional network underlying the T_Eff-2_ state.

**Figure 4:**
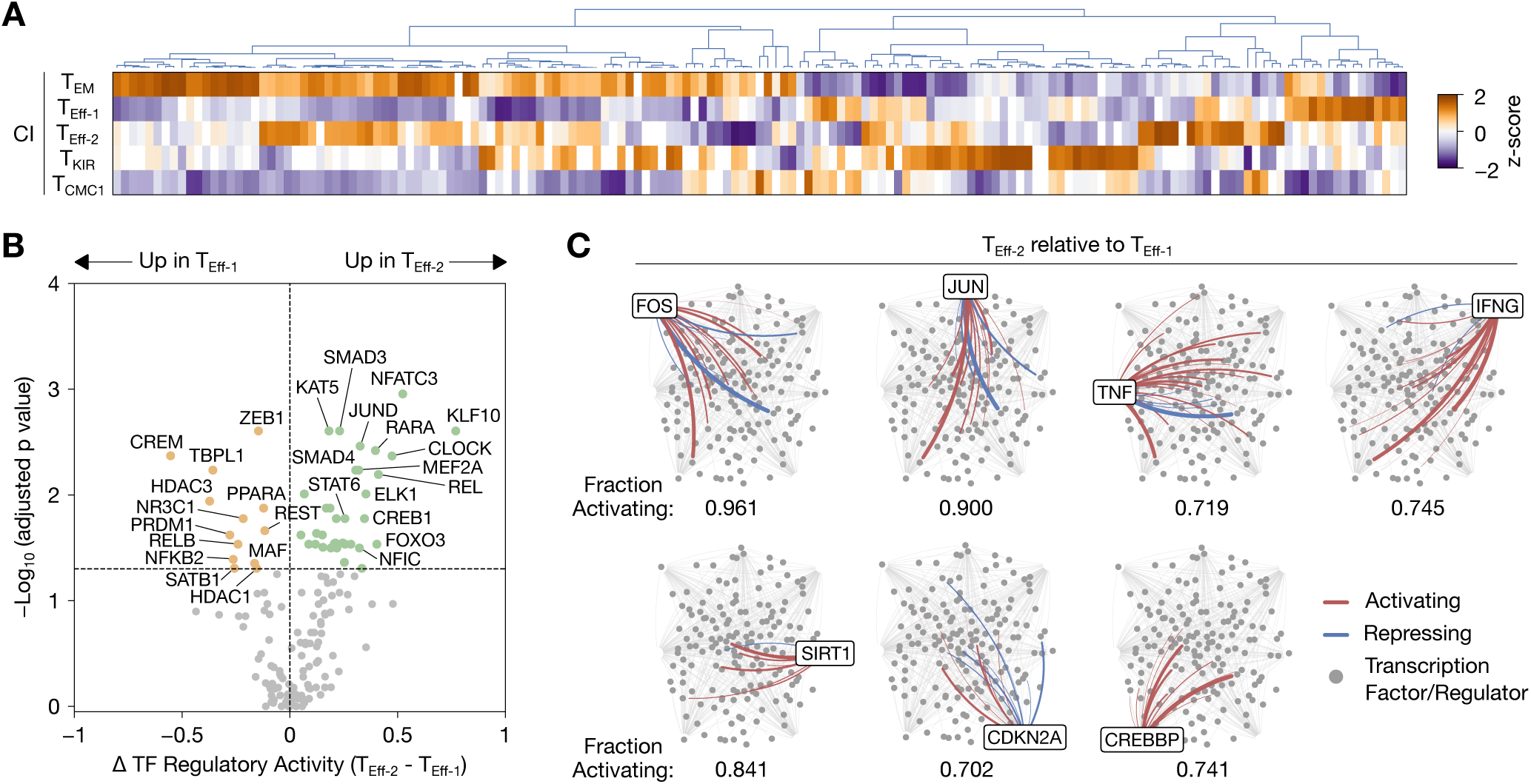
Gene regulatory networks support both activation and metabolic restraint. **(A)** Univariate linear modeling (ULM) on a per donor basis of the CollecTRI transcription factor network in CD8^+^ T cell subsets. Transcription factors are arranged by hierarchical clustering. **(B)** Differential regulatory activity of transcription factors in T_Eff-2_ relative to T_Eff-1_. **(C)** Network plots for top candidate transcription factor gene targets identified by delta-ULM-weighted GRN target analysis comparing T_Eff-2_ versus T_Eff-1_. The full network connectivity is shown in gray. Each grey circle represents a transcription factor/regulator in the analysis. Those connections with significant signed regulator-target edges are highlighted (red, activating; blue, repressing). The full network connectivity is shown in gray. Line weight shows the strength of effect as determined by ULM in **(A)**. The overall fraction of significant regulator-target gene edges supporting activation of the listed gene, weighted by delta-ULM strength, is shown. Statistical testing for **(B)** was performed using the Mann-Whitney U test with Benjamini-Hochberg adjustment. TF, transcription factor.

### T_Eff-2_ occupies a terminal position within the CD8^+^ T cell compartment

The transcriptional features of T_Eff-2_ suggested a cell state that diverges from canonical effector differentiation, prompting us to investigate its position within the CD8^+^ T cell differentiation hierarchy. RNA velocity analysis using VeloVI (34) identified directional flow from T_EM_ toward both T_Eff-1_ and T_Eff-2_, with additional streams from T_Eff-1_ toward T_Eff-2_, suggesting T_Eff-2_ represents a more terminal state (Fig. 5A, Supplemental Fig. 3). To quantify differentiation dynamics, we computed a CellRank2 (35) PseudotimeKernel integrating Palantir pseudotime (36) with transcriptomic similarity-based neighbor graph and simulated random walks from T_N_ cells (Fig. 5B). Random walks preferentially terminated in T_Eff-2_ alongside T_Eff-1_, T_CMC1_, and T_KIR_, consistent with T_Eff-2_ representing one of several terminal circulating CD8^+^ T cell fates. Palantir pseudotime corroborated this positioning, with T_Eff-2_ occupying a more terminal pseudotime distribution than T_Eff-1_ (Fig. 5C).

**Figure 5:**
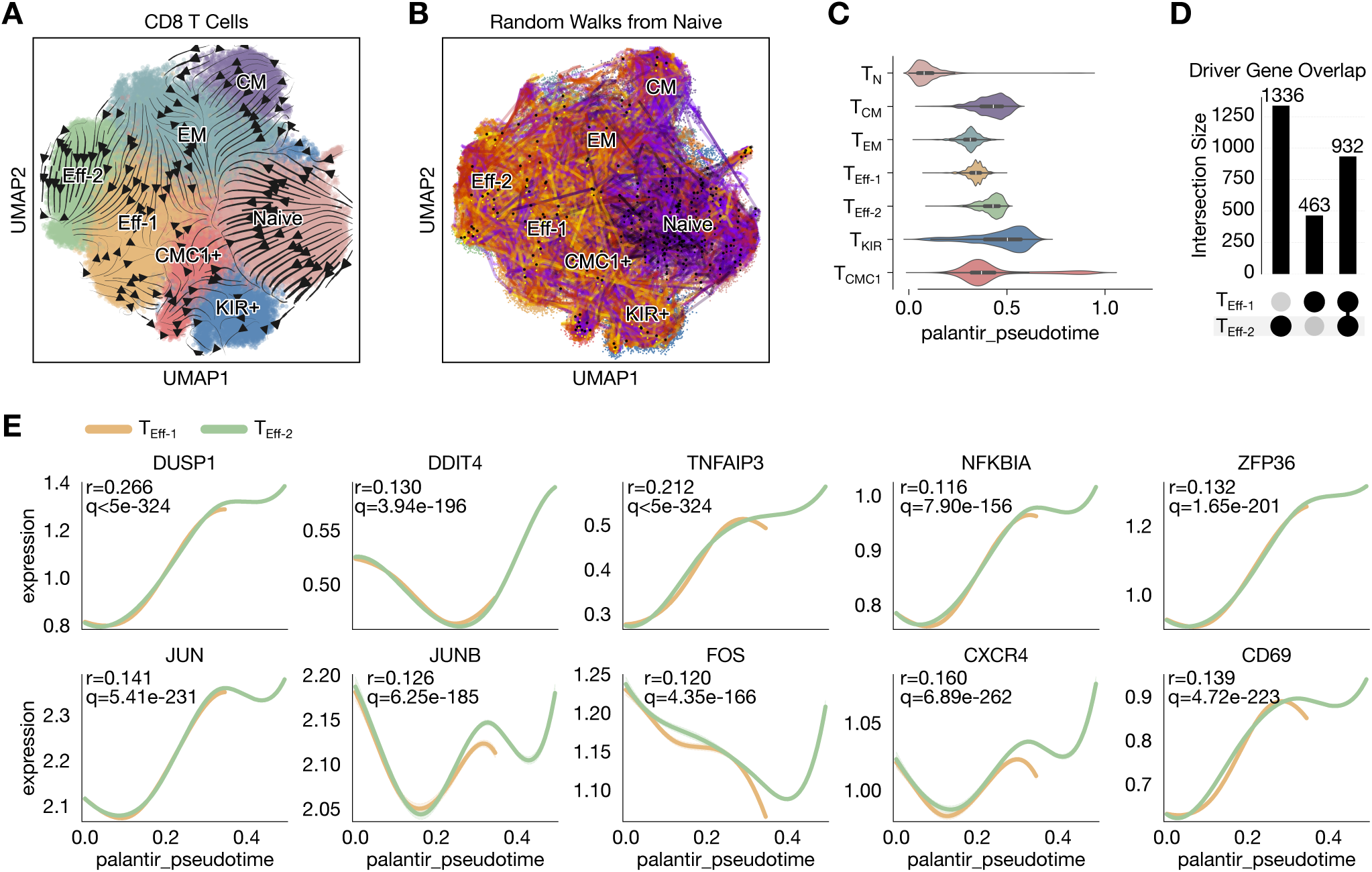
RNA velocity and pseudotime analyses identify T_Eff-2_ as a terminal fate. **(A)** UMAP visualization of the RNA velocity stream computed by VeloVI. **(B)** Random walks initiated from T_N_ on the CellRank2 PseudotimeKernel projected onto the UMAP embedding. Black and yellow dots indicate random walk start and end points, respectively. **(C)** Palantir pseudotime distributions across CD8^+^ T cell clusters, rooted in T_N_. **(D)** UpSet plot of significant lineage driver genes (q < 0.05) for T_Eff-2_ and T_Eff-1_ fates identified by CellRank2 fate probability-weighted generalized additive model regression using MAGIC-imputed expression values. **(E)** Gene expression trends for selected top T_Eff-2_ lineage driver genes along Palantir pseudotime modeled by generalized additive model using MAGIC-imputed expression values for T_Eff-2_ and T_Eff-1_ lineages.

To identify gene programs driving commitment to the T_Eff-2_ fate, we performed CellRank2 fate probability-weighted lineage driver analysis comparing T_Eff-2_ and T_Eff-1_ (Fig. 5D). T_Eff-2_ was associated with 1,336 significant (q < 0.05) fate-specific driver genes versus 463 for T_Eff-1_, indicating that commitment to the T_Eff-2_ fate is driven by a substantially more extensive lineage-specific transcriptional program. Gene expression trends along pseudotime showed progressive enrichment of genes reflecting metabolic restraint and active inflammatory regulation in T_Eff-2_ relative to the T_Eff-1_ lineage, including the mTORC1 suppressor *DDIT4*, the MAPK inhibitor *DUSP1*, the NF-κB negative feedback regulators *TNFAIP3* and *NFKBIA*, and the inflammatory mRNA decay factor *ZFP36* (Fig. 5E). Genes reflecting an activated effector state showed similar enrichment, including the AP-1 transcription factors *JUN*, *JUNB*, and *FOS*, the early activation marker *CD69*, and the chemokine receptor *CXCR4*. Together, these data support T_Eff-2_ as a terminal effector state.

### Transcriptomic resolution is required to identify CD8^+^ T_Eff-2_

We next sought to identify CD8^+^ T_Eff-2_ using classical protein-based cytometric methods. Since T_Eff-2_ cells are transcriptomically terminal effectors, we considered if they represented T effector memory cells re-expressing CD45RA (T_EMRA_) cells in CI. We first assessed whether flow cytometry (FC)-defined T_EMRA_ frequency could resolve T_Eff-2_. Despite T_Eff-2_ being present at appreciable frequencies only in CI, T_EMRA_ frequency did not differ significantly between HC and CI (Fig. 6A, B), suggesting that T_Eff-2_ expansion is obscured within the broader T_EMRA_ compartment. FC-defined T_N_ frequency correlated strongly with scRNA-seq T_N_ proportions, validating this cross-platform comparative approach (Fig. 6C). FC-defined T_EMRA_ frequency correlated only moderately with scRNA-seq T_Eff-1_ and T_Eff-2_ proportions individually but more strongly with their combined proportion, consistent with both populations substantially overlapping with FC-defined T_EMRA_ cells but being insufficiently distinct to discriminate one from the other.

**Figure 6:**
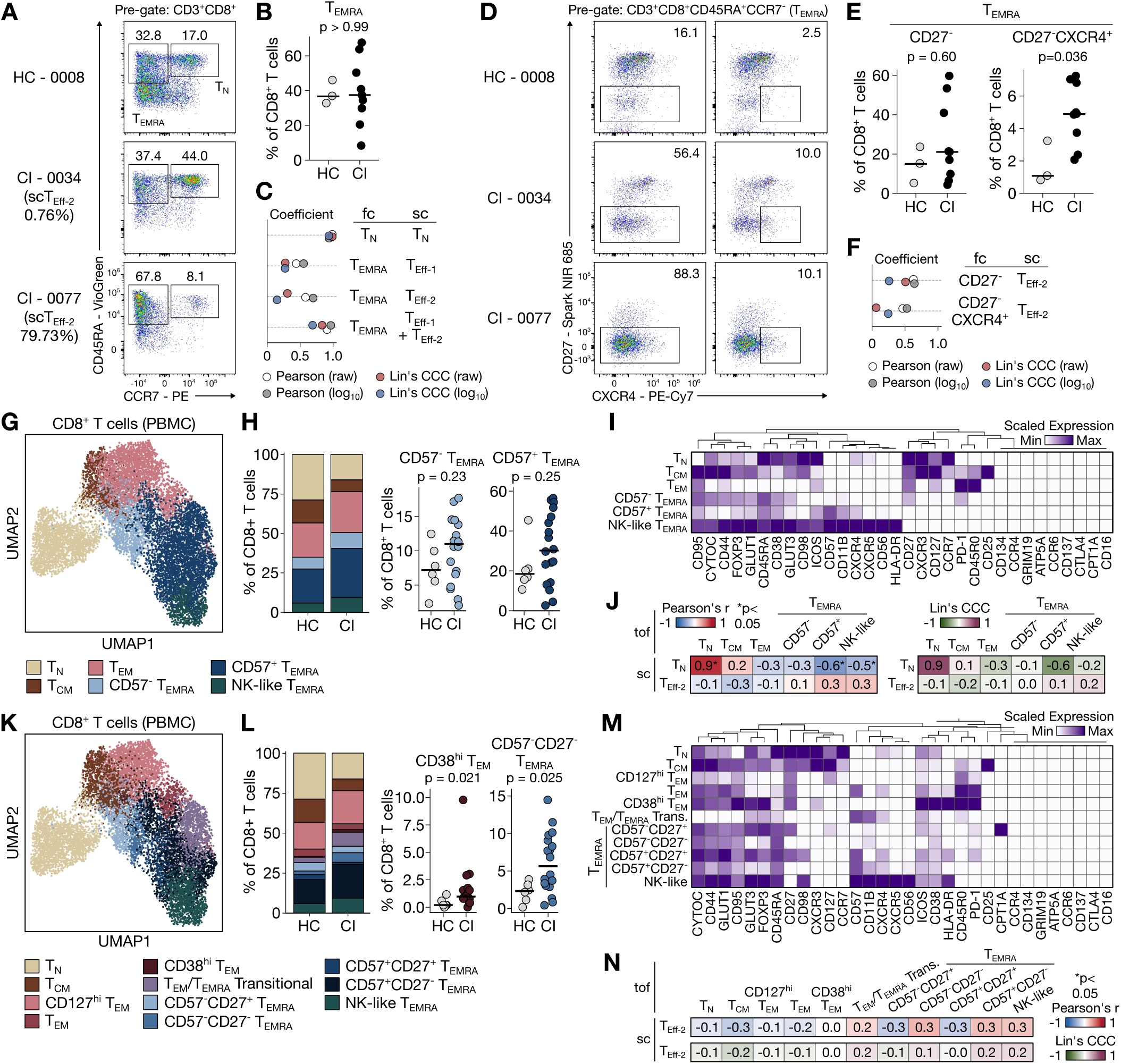
CD8^+^ T_Eff-2_ is not resolved at the protein level by conventional flow cytometry or mass cytometry. **(A)** Representative flow cytometry plots of CD45RA and CCR7 expression in CD3^+^CD8^+^ T cells from a HC participant, a CI patient with low scRNA-seq T_Eff-2_ frequency (0.76%), and a CI patient with high scRNA-seq T_Eff-2_ frequency (79.73%). **(B)** Pooled frequency of CD8^+^ T_EMRA_ (CD3^+^CD8^+^CD45RA^+^CCR7^-^) identified by flow cytometry as a proportion of all CD8^+^ T cells in HC and CI. **(C)** Pearson correlation coefficients and Lin’s concordance correlation coefficients (CCC) of CD8^+^ T_N_ and T_EMRA_ frequencies identified by flow cytometry compared to corresponding scRNA-seq cluster proportions, shown for raw and log_10_-transformed data. **(D)** Representative flow cytometry plots of CXCR4 and CD27 expression on CD8^+^ T_EMRA_ from the same samples as in **(A)** gated for CD27^-^ T_EMRA_ (left) and CD27^-^CXCR4^+^ T_EMRA_ (right). **(E)** Pooled frequencies of CD27^-^ T_EMRA_ (left) and CD27^-^CXCR4^+^ T_EMRA_ (right) as a proportion of all CD8^+^ T cells in HC and CI. **(F)** Pearson correlation coefficients and Lin’s CCC of CD27^-^ and CD27^-^CXCR4^+^ T_EMRA_ frequencies compared to scRNA-seq T_Eff-2_ proportions, shown for raw and log_10_-transformed data. **(G)** UMAP visualization of CD8^+^ T cells from PBMCs of HC and CI patients measured by mass cytometry (CyTOF), annotated by cluster. **(H)** Mean frequency of each CyTOF CD8^+^ T cell cluster among all CD8^+^ T cells in HC and CI patients (left) and per participant frequencies of CD57^-^ T_EMRA_ and CD57^+^ T_EMRA_ clusters (right). **(I)** Heatmap of median scaled expression of clustering markers across CD8^+^ T cell clusters. **(J)** Heatmap of Pearson correlation coefficients and Lin’s CCC of scRNA-seq T_N_ and T_Eff-2_ cluster proportions compared to all CyTOF cluster proportions. **(K)** UMAP visualization of CD8^+^ T cells from PBMCs of HC and CI patients measured by CyTOF at higher clustering resolution compared to **(G)**, annotated by cluster. **(L)** Mean frequency of each higher-resolution CyTOF CD8^+^ T cell cluster among all CD8^+^ T cells in HC and CI patients (left) and per participant frequencies of CD38^hi^ T_EM_ and CD57^-^CD27^-^ T_EMRA_ clusters (right). **(M)** Heatmap of median scaled expression of clustering markers across higher-resolution CD8^+^ T cell clusters. **(N)** Heatmap of Pearson correlation coefficients and Lin’s CCC of scRNA-seq T_Eff-2_ cluster proportions compared to all high-resolution CyTOF cluster proportions. Statistical testing was performed using the Mann-Whitney U test **(B, E, H, L)** and the Pearson correlation coefficient with two-sided p value with significant p values < 0.05 denoted **(C, F, J, N)**. Lin’s concordance correlation coefficient were also reported **(C, F, J, N)**. Raw (untransformed) proportions **(C, F, J, N)** and log_10_-transformed proportions **(C, F)** were used to calculate statistics. Each datapoint **(B, E, H, L)** represents an individual research participant. sc = scRNA-seq; tof = CyTOF.

Relatively few cell surface markers are differentially expressed between T_Eff-1_ and T_Eff-2_. Given that *CXCR4* (log_2_fc = 0.55, p_adj_ = 0.00000068) and *CD27* (log_2_fc = −1.36, p_adj_ = 0.00048) transcripts are among those differentially expressed in T_Eff-2_, we next attempted to enrich for T_Eff-2_ within the T_EMRA_ gate using these markers. CD27^-^ T_EMRA_ frequency did not differ between groups, while CD27^-^CXCR4^+^ T_EMRA_ frequency was significantly higher in CI than HC (Fig. 6D, E). However, neither population correlated with scRNA-seq T_Eff-2_ proportions (Fig. 6F). Additionally, among CI patients, CD27^-^CXCR4^+^ T_EMRA_ frequencies in the most abundant samples remained less than 10% of CD8^+^ T cells, whereas T_Eff-2_ frequencies in the paired scRNA-seq samples reached 79.73% in one sample (Fig. 6A, E).

Since T_Eff-2_ were identified through clustering in high-dimensional transcriptomic space, we reasoned that an analogous high-dimensional protein-level approach might better capture this population. We performed mass cytometry (CyTOF) using a T-cell focused panel of 43 markers on 24 PBMC samples (6 HC, 18 CI), clustering CD8^+^ T cells to recapitulate the scRNA-seq defined populations and identifying 6 clusters as the closest protein-level analogs (Fig. 6G). Among these samples, 22 (6 HC, 16 CI) were matched aliquots from the same blood collections used for scRNA-seq allowing for robust frequency comparisons. Neither CD57^-^ T_EMRA_ nor CD57^+^ T_EMRA_ proportions differed significantly between HC and CI (Fig. 6H). Median scaled expression of markers across clusters is shown (Fig. 6I). To assess whether any CyTOF cluster captured T_Eff-2_, we correlated scRNA-seq T_N_ and T_Eff-2_ proportions with all CyTOF cluster proportions (Fig. 6J, Supplemental Fig. 5A). scRNA-seq T_N_ proportions correlated strongly with CyTOF T_N_ proportions (Pearson’s r=0.94, Lin’s CCC=0.91), confirming that CyTOF clustering can recapitulate scRNA-seq-defined populations. However, no CyTOF cluster showed meaningful correlation with scRNA-seq T_Eff-2_ proportions.

To determine if T_Eff-2_ had been merged with other populations at this clustering resolution, we repeated the analysis using higher resolution clustering (Fig. 6K). Only CD38^hi^ T_EM_ and CD57^-^CD27^-^ T_EMRA_ trended toward higher proportions in CI than HC (Fig. 6L). Median scaled expression of markers across clusters is shown (Fig. 6M). Correlating scRNA-seq T_Eff-2_ proportions with higher-resolution CyTOF cluster proportions revealed no meaningful association (Fig. 6N, Supplemental Fig. 5B). We reasoned that in sample V-0055, where T_Eff-2_ comprised 72.79% of the CD8^+^ T cells by scRNA-seq, the strongest CD8^+^ T cell protein expression signals would largely reflect T_Eff-2_. We selected the 10 markers most differentially expressed in total CD8^+^ T cells in V-0055 and reclustered the full CyTOF dataset but still observed no meaningful correlation with scRNA-seq T_Eff-2_ proportions (Supplemental Fig. 5C-E). Restricting analysis to CD8^+^CCR7^-^ T cells to reduce T_N_ and T_CM_ influence on clustering similarly failed to identify a correlated cluster (Supplemental Fig. 5F-H).

Our prior clustering approaches were unsupervised and therefore agnostic to scRNA-seq T_Eff-2_ frequency. We next applied CITRUS (37), which combines hierarchical clustering with supervised learning to identify subpopulations whose abundance associates with a sample-level endpoint, using scRNA-seq T_Eff-2_ proportions as the outcome. No clusters were positively associated with T_Eff-2_ frequency at FDR < 0.05 in either raw or logit-transformed data (Supplemental Table 6). Collectively, the T_Eff-2_ state was poorly resolved by protein cytometry across multiple clustering and feature-selection strategies, indicating effective camouflaging within standard immunophenotypic space.

### The T_Eff-2_ transcriptional program shares features with lung tissue-resident memory CD8^+^ T cells in pneumonia

The association of T_Eff-2_ with severe hypoxemic respiratory failure prompted us to ask whether circulating T_Eff-2_ shares transcriptomic features with CD8^+^ T cell populations in the inflamed lung. To test this, we utilized a publicly available scRNA-seq dataset (GSE167118) (38) of sorted CD3^+^ bronchoalveolar lavage (BAL) cells from mechanically ventilated patients with severe SARS-CoV-2 pneumonia (n=7) or bacterial pneumonia (n=4). We restricted our analysis to CD8^+^ T cells, identifying 5 transcriptionally distinct populations including effector memory T cells (T_EM_), T cells with high interferon-stimulated gene expression (T_ISG_), cytotoxic effector T cells (T_CyTox_), tissue-resident memory T cells (T_RM_), and mucosal-associated invariant T cells (MAIT) (Fig. 7A-C).

**Figure 7:**
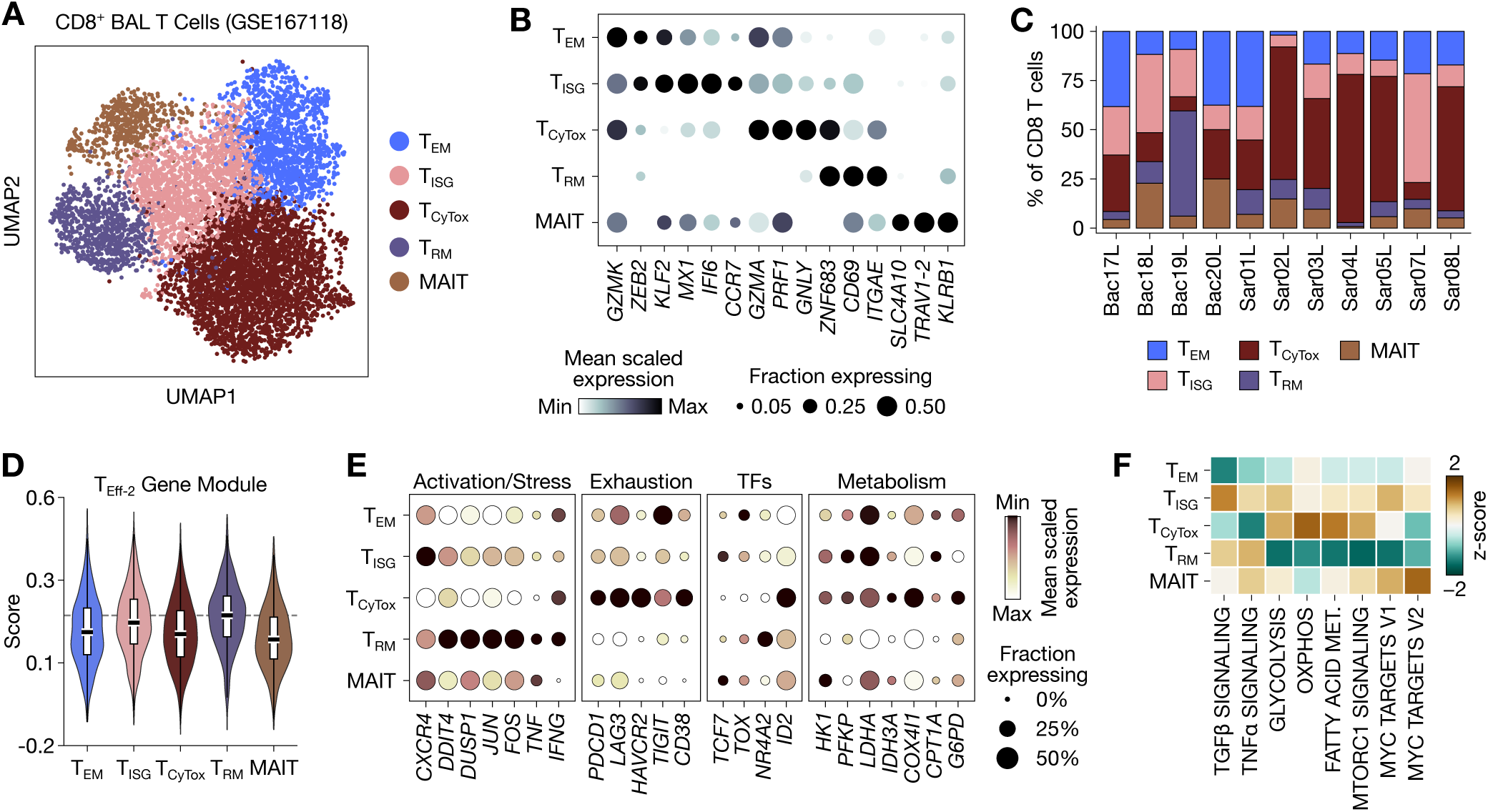
Circulating CD8^+^ T_Eff-2_ share transcriptomic features with bronchoalveolar lavage (BAL) tissue resident memory CD8^+^ T cells. **(A)** UMAP visualization of CD8^+^ T cells from BAL of bacterial and SARS-CoV-2 pneumonia patients (GSE167118) annotated by cell cluster. **(B)** Selected BAL CD8^+^ T cell subset-defining genes for each cell cluster in **(A)**. **(C)** Frequencies of each BAL CD8^+^ T cell subset among all CD8^+^ T cells in the BAL, shown per patient. Samples annotated with ‘Bac’ are bacterial pneumonia; samples annotated with ‘Sar’ are SARS-CoV-2 pneumonia. **(D)** Circulating CD8^+^ T_Eff-2_ gene module score among BAL CD8^+^ T cell subsets. The dashed line represents the median T_RM_ score. **(E)** Normalized scaled gene expression of selected circulating CD8^+^ T_Eff-2_ functional and regulatory genes in BAL CD8^+^ T cell subsets. **(F)** Univariate linear modeling (ULM) on a per cell basis of Hallmark pathways in BAL CD8^+^ T cell subsets.

To assess transcriptomic overlap between circulating T_Eff-2_ and BAL CD8^+^ T cell subsets, we computed a T_Eff-2_ gene module score (Fig. 7D, Supplemental Table 7), finding that T_RM_ cells had the highest T_Eff-2_ module score across BAL CD8^+^ T cell subsets. Consistent with this, T_RM_ cells had elevated expression of T_Eff-2_ signature genes including *DDIT4*, *DUSP1*, *JUN*, *FOS*, *TNF*, and *IFNG*, alongside reduced expression of glycolytic and mitochondrial metabolic genes (*HK1*, *LDHA*, *IDH3A*, *COX4I1*, and *CPT1A)*, relative to other BAL CD8^+^ T cell subsets (Fig. 7E). At the pathway level, T_RM_ cells similarly showed positive enrichment of TGFβ signaling consistent with tissue residence (Fig. 7F). T_RM_ also exhibited TNFα signaling enrichment alongside broad downregulation of metabolic pathways including glycolysis, oxidative phosphorylation, fatty acid metabolism, and mTORC1 signaling relative to other BAL CD8^+^ T cell subsets, mirroring the hypometabolic and inflammatory transcriptional signature of circulating T_Eff-2_ (Fig. 7F).

### T_Eff-2_ frequency and phenotype were validated in an independent external sepsis cohort

To validate the T_Eff-2_ population in an independent cohort, we utilized a publicly available scRNA-seq PBMC dataset (GSE279452) (29) from adult critically ill patients with sepsis (CI-Sep; n=91) and healthy controls (HC; n=7). We clustered CD8^+^ T cells from this dataset, identifying the T_Eff-2_ state, which was only present in substantial proportions in sepsis patients (Fig. 8A-C).

**Figure 8:**
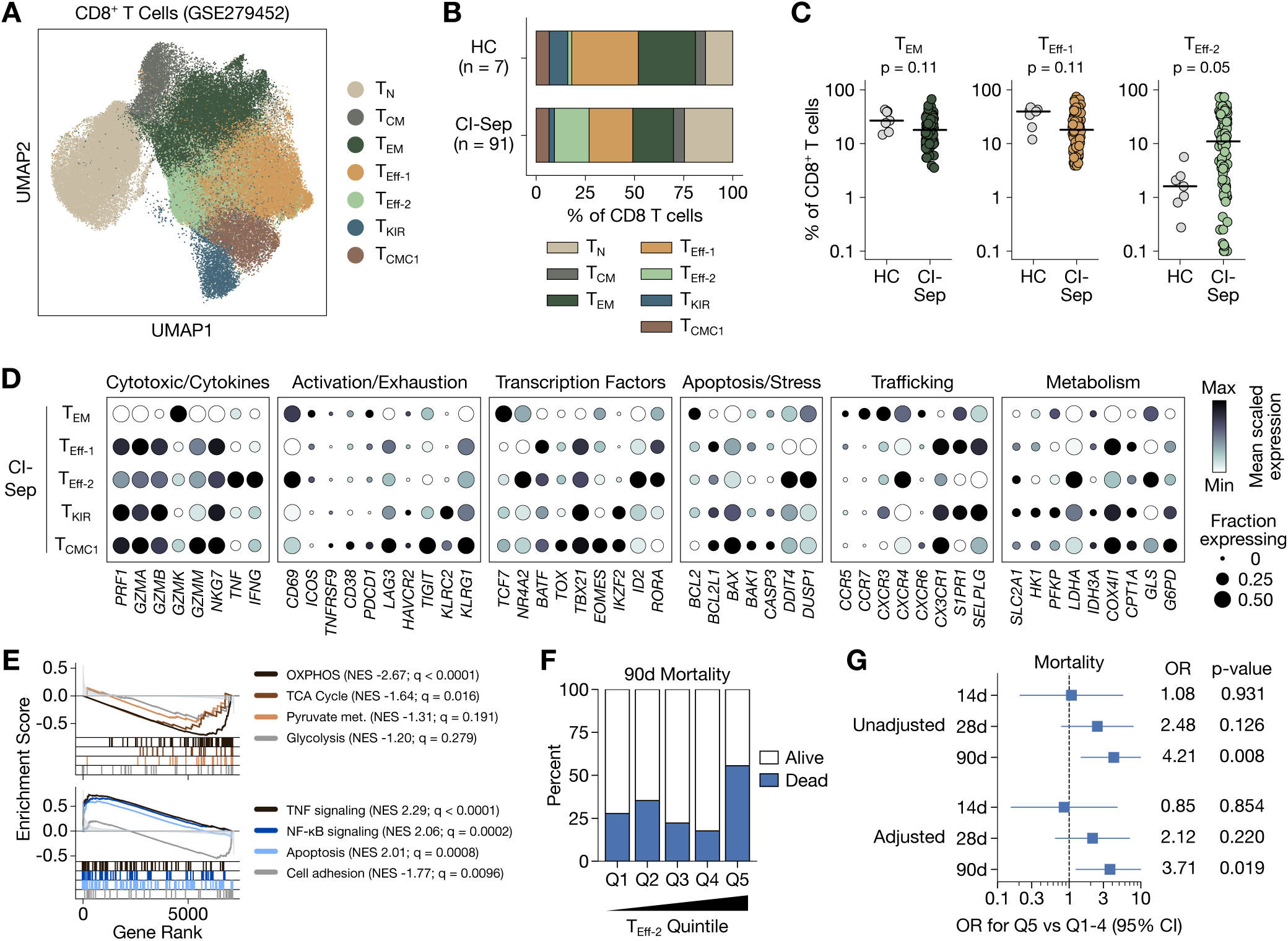
Validation of CD8^+^ T_Eff-2_ frequency and phenotype in an external sepsis cohort. **(A)** UMAP visualization of CD8^+^ T cells from HC and CI patients with sepsis (CI-Sep, GSE279452) annotated by cell cluster. **(B)** Mean proportion of each T cell cluster among all CD8^+^ T cells in HC and CI-Sep patients. **(C)** Per participant frequencies of effector CD8^+^ T cell clusters among all CD8^+^ T cells in HC and CI-Sep patients. **(D)** Normalized scaled gene expression of selected functional and regulatory genes in CI-Sep CD8^+^ T cell subsets. **(E)** Gene set enrichment analysis (GSEA) of selected KEGG pathways comparing CI-Sep T_Eff-2_ to CI-Sep T_Eff-1_. **(F)** Mortality at 90 days by quintiles of T_Eff-2_ proportion. **(G)** Unadjusted and age- and sex-adjusted logistic regressions of the highest frequency quintile (Q5) compared to all other quintiles (Q1-4) for 14-, 28-, and 90-day mortality. Each datapoint **(C)** represents an individual research participant. Statistical analysis was performed using the Empirical Bayes moderated T-test **(C)** and the permutation-based Kolmogorov–Smirnov statistic with Benjamini–Hochberg correction **(E)**. NES, normalized enrichment score; OR, odds ratio.

Normalized gene expression of T_Eff-2_ in this external cohort recapitulated key features of its activated and stressed identity, with elevated expression of *TNF*, *IFNG*, *CD69*, *NR4A2*, *ID2*, *RORA*, *DDIT4*, *DUSP1*, and *CXCR4* and minimal expression of *TIGIT* (Fig. 8D). The metabolic signature was largely recapitulated, with reduced *HK1*, *PFKP*, *IDH3A*, and *COX4I1* expression relative to other effector lineages. *LDHA* was an exception, showing high expression in T_Eff-2_ in this validation cohort. Gene set enrichment analysis comparing T_Eff-2_ to T_Eff-1_ showed significant enrichment of TNF and NF-κB signaling and significant depletion or a trend toward reduction of oxidative phosphorylation, TCA cycle, glycolysis, and pyruvate metabolism pathways (Fig. 8E). These findings are consistent with the inflammatory-conditioned and hypometabolic transcriptional identity of circulating T_Eff-2_ in our cohort.

We used the 32 sepsis samples in this validation dataset with paired Cellular Indexing of Transcriptomes and Epitopes by Sequencing (CITE-seq) data to search for additional protein marker(s) that could enable flow cytometric labeling of T_Eff-2_. Among putative lineage-defining markers (median expression above isotype control in ≥50% of cells in at least one CD8^+^ T cell cluster), T_Eff-2_ uniquely showed elevated CD101 expression (Supplemental Fig. 6A–D). CD41 showed similarly elevated ADT signal but was excluded as a platelet-specific marker. Notably, *CD101* transcript abundance was quite low across the CD8^+^ T cell clusters, making CD101 a candidate marker not identified by our previous analyses that were informed by RNA-based differential expression (Supplemental Fig. 6E). Flow cytometry incorporating CD101 showed no positive correlation between CD101^+^ CD8^+^ T cell frequency and T_Eff-2_ proportions in matched flow and scRNA-seq samples, including when analyses were restricted to effector lineage cells (CCR7^-^ CD8+ T cells) or combined with CXCR4 or CD27 expression (Supplemental Fig. 6F,G). In fact, gating for CD101 had a trend for negative correlations with T_Eff-2_ across most analyses (Supplemental Fig. 6G). This approach again failed to resolve T_Eff-2_ at the protein level.

To test whether T_Eff-2_ frequency corresponded to adverse clinical outcomes in this independent cohort, we examined mortality across quintiles of T_Eff-2_ proportion. Patient-level PaO_2_/FiO_2_ data were unavailable, precluding direct validation of the hypoxemic respiratory failure association. However, available 14-, 28-, and 90-day mortality data enabled us to validate whether T_Eff-2_ frequency was associated with mortality. Given the skewed distribution of T_Eff-2_ frequencies with marked enrichment in the upper quintile (Supplemental Fig. 7), we compared the highest quintile to the lower four quintiles using logistic regression. Patients in the highest T_Eff-2_ frequency quintile had increased 90-day mortality compared to patients in the lower four quintiles (Fig. 8F), an association that remained significant even after adjustment for age and sex (Fig. 8G). This mortality association strengthened over time from hospital admission, with no association at day 14, a non-significant trend at day 28, and a significant effect by day 90. Collectively, these findings validate T_Eff-2_ as a critical illness-expanded CD8^+^ T effector state with mixed inflammatory and hypometabolic features associated with adverse clinical outcomes.

## Discussion

Herein, we identify T_Eff-2_, a novel CD8^+^ effector subset that is expanded in critical illness from both septic and non-septic etiologies. T_Eff-2_ frequency associated selectively with hypoxemic respiratory failure and mortality, and its transcriptional identity and clinical associations were reproduced in an independent external sepsis cohort. Trajectory analyses position T_Eff-2_ as a terminal effector fate driven by repressors of mTOR and MAPK, and its transcriptional program shares features with alveolar CD8^+^ T_RM_ cells in pneumonia.

The expansion of T_Eff-2_ across both non-septic and septic ICU patients suggests its induction reflects shared physiological features of critical illness rather than pathogen-driven immune activation. Its transcriptional program, characterized by mTORC1 suppression, AP-1 activation, MAPK and NF-κB negative feedback, and broad metabolic restraint alongside preserved inflammatory cytokine expression, is consistent with a cellular response to environmental stress. Recent work demonstrated that murine CD8^+^ T cells engage an integrated stress response (ISR) under nutrient stress, inducing a transcriptional program marked by repression of mTORC1 pathway activity analogous to what we observed in T_Eff-2_ (39). Critical illness imposes hypoperfusion, tissue hypoxia, and nutrient dysregulation, each sufficient to activate the ISR (40–43). These observations raise the possibility that T_Eff-2_ reflects an ISR-conditioned state, though further mechanistic work in human critical illness is needed. Aligned with this possibility, prolonged ISR activation can drive cellular senescence (44), and our T_Eff-2_ gene regulatory analysis demonstrated convergent activation of *CDKN2A* (encoding the senescence regulator p16) alongside elevated *FOXO3* activity, which both associate with a senescence-like state.

The ontogeny of T_Eff-2_ remains not fully resolved. The selective association of T_Eff-2_ with hypoxemic respiratory failure may reflect a relationship with pulmonary immune responses. T_Eff-2_ cells may arise independently through shared environmental stressors or represent ex-T_RM_ cells that have re-entered circulation, a possibility supported by studies demonstrating CD8^+^ T_RM_ cell egress under inflammatory conditions (45–47). Consistent with the latter, the T_Eff-2_ transcriptional program was partially recapitulated in BAL T_RM_ cells from pneumonia patients. Further work is needed to delineate the origins of this state, as well as whether a similar state exists in non-critical illness or chronic inflammatory disease.

The study from which the external validation cohort was derived (29) also identified CD8^+^ T cell heterogeneity in sepsis. In that analysis, cluster T_CD8_c05-GNLY-CCL4^hi^ notably shares some features with T_Eff-2_, including preserved cytotoxic gene expression, upregulated pro-inflammatory mediators, and an association with mortality. However, the transcriptional profiles do not fully align. Specifically, the defining marker of their cluster, *CCL4*, was not significantly differentially expressed in T_Eff-2_ vs T_Eff-1_ in our data (log_2_FC 0.105, p_adj_ 0.996; Supplemental Table 2), and key T_Eff-2_-defining genes including *DDIT4* and *DUSP1* were not significantly enriched in their cluster. These findings indicate partial transcriptional overlap but support T_Eff-2_ as a distinct stress-adapted effector program not fully captured in this prior analysis.

Despite extensive investigation using flow cytometry, mass cytometry, and CITE-seq-guided marker discovery, T_Eff-2_ could not be resolved at the protein level. We propose that T_Eff-2_ exemplifies what we term as a “latent transcriptional state” defined by a highly reproducible transcriptomic program, but whose distinguishing features are not recognizable by standard protein-based cytometric phenotyping approaches. Transcriptomic clusters are defined by covariance across hundreds of genes, making mapping into lower dimensional protein-based cytometry inherently reductive. Additionally, post-translational regulation and mRNA-protein decoupling may further obscure the correspondence between transcriptional and protein-level identity. Furthermore, commercially available flow and mass cytometry reagents provide broad coverage of some protein classes (e.g., cluster of differentiation markers) but sparsely cover others (e.g., metabolic proteins and transcription factors), limiting the ability of protein-based cytometry to capture all relevant cellular states.

Critical illness syndromes including sepsis and acute respiratory distress syndrome (ARDS) are highly heterogenous, a feature believed to have contributed to the failure of major ICU clinical trials. Latent classes in these syndromes, such as the hypo- and hyperinflammatory states defined largely by plasma biomarkers (48) and the sepsis response signatures (SRS1/SRS2) derived from bulk leukocyte transcriptomics (24) have been validated in multiple cohorts. In retrospective analyses of randomized ICU trials, these latent classes identify patient subgroups that may have benefited from therapies that showed no overall effect in the original studies (49–52). Our work extends this framework beyond plasma biomarkers and bulk RNA-sequencing by identifying a reproducible, single-cell-defined transcriptional state that associates with clinically significant outcomes but which is not easily resolved by standard cytometry-based immunophenotyping. Major global funding agencies have invested in consortia to more precisely phenotype and translate discoveries in critical illness including the ARDS, Pneumonia, and Sepsis (APS) Consortium (NIH/United States), BEATSEPSIS (Horizon Europe/European Union), and the Chinese Multi-omics Advances in Sepsis (CMAISE) cohort (China). Our data underscore the need to include scRNA-seq and related single-cell transcriptomic approaches alongside traditional plasma biomarker and bulk RNA-seq to avoid missing clinically meaningful axes of immune dysfunction.

This study has several limitations. Our primary cohort was from a single institution, though results were validated externally. Enrollment was restricted to medical and surgical ICU patients, and generalizability to populations such as trauma patients is unknown. Many functional characterizations were not possible given our inability to resolve T_Eff-2_ at the protein level, and its ontogeny and relationship to BAL T_RM_ cells remains unresolved.

Despite these limitations, our study characterizes CD8^+^ T cell heterogeneity in human critical illness at single-cell resolution across septic and non-septic etiologies, with external validation in an independent sepsis cohort. Convergent transcriptional, regulatory, and trajectory evidence supports T_Eff-2_ as a reproducible, clinically significant effector state whose identification required single-cell transcriptomic resolution. These findings advance our understanding of CD8^+^ T cell biology in human critical illness and highlight the necessity of single-cell transcriptomic approaches for resolving clinically meaningful immune substructure. Such approaches will be valuable for identifying therapeutic targets and developing precision strategies to restore effective immunity in critical illness.

## Methods

### Sex as a biological variable

This study enrolled and analyzed samples from male and female participants (Table 1). Biological sex was included as a covariate in the scVI integration model and in external validation mortality regression analyses. Biological sex was not examined as a primary variable of interest.

### Human subjects and clinical metadata

Critically ill research participants were enrolled from adults admitted to the medical and surgical intensive care units (ICUs) at Vanderbilt University Medical Center from 2022 to 2023 as part of the Sepsis ClinicAl Resource And Biorepository (SCARAB) as previously described (30). Sepsis status was assigned using modified Sepsis-3 criteria by an ICU physician as previously described (30). Non-acutely ill, non-hospitalized adults were recruited as healthy controls as previously described (30).

### PBMC Isolation and Storage

SepMate-50 tubes (STEMCELL Technologies, cat. no. 85450) were loaded with 15 ml Lymphoprep (STEMCELL Technologies, cat. no. 18061), and fresh peripheral blood diluted 1:2 in PBS was layered on top. Following centrifugation at 1,200g for 15 min at 20°C, the PBMC-containing layer was collected, pelleted at 400g for 10 min at 20°C, and resuspended in ACK lysis buffer (Gibco, cat. no. A1049201) for 7 min. Cells were then washed twice at 300g for 10 min at 20°C and passed through a 70-μm strainer. PBMCs were resuspended at 2 million cells per mL in Bambanker Cell Freezing Medium (GC LYMPHOTEC, cat. no. BB02), aliquoted, and frozen in a CoolCell FTS30 container (Corning, cat. no. 432006) at −80°C before transferring to liquid nitrogen vapor-phase storage within 72 h.

### PBMC Cryorecovery

PBMC aliquots were thawed at 37°C for 3 min and diluted 1:5 into HPLM (Gibco, cat. no. A4899101) supplemented with 10% dialyzed fetal bovine serum (dFBS; Sigma-Aldrich, cat. no. F0392) and 1% penicillin-streptomycin (Gibco, cat. no. 15140122). Cells were pelleted at 300g for 5 min at 20°C and washed twice in the same medium, with the final wash performed through a 35-μm cell strainer cap. PBMCs were resuspended in HPLM with 10% dFBS and 1% penicillin-streptomycin for immediate use.

### Single-cell RNA sequencing library preparation and data pre-processing

Single-cell RNA sequencing data were generated as part of a previously published PBMC dataset (30). Briefly, cryopreserved PBMCs were thawed, washed, filtered, and loaded for 5′ single-cell RNA sequencing using the 10x Genomics Chromium Next GEM Single Cell 5′ HT Kit v2 platform. Libraries were sequenced on an Illumina NovaSeq 6000, and raw sequencing data were aligned to the human GRCh38-2020-A reference genome using Cell Ranger (v7.1.0).

The parent PBMC dataset was processed as described, including ambient RNA and artifact correction (CellBender v0.3.0), cell-level quality control (>20% mitochondrial content or <400 or >5,000 detected features excluded), doublet removal (scDblFinder v1.13.10), and batch-aware latent space integration (scVI v1.0.4). Within this parent PBMC dataset, cells annotated as CD8 T cells (celltype_coarse = “CD8 T”) were retained to generate a new AnnData object for downstream CD8-specific analyses. Innate-like mucosal-associated invariant T (MAIT) cells, which often express the co-receptor CD8, were not included in this object.

### CD8^+^ T cell integration, clustering, and subset annotation

The CD8^+^ T cell object was reprocessed independently from the parent PBMC object using Scanpy (v1.9.5). Genes with fewer than 3 total counts were removed. Highly variable genes were selected (flavor = “seurat_v3”, n_top_genes = 10,000) and used for downstream scVI integration. The scVI model was specified with n_hidden = 128, n_latent = 10, n_layers = 1, dispersion = “gene”, and gene_likelihood = “zinb”. Categorical (biological sex) and continuous (mitochondrial read percentage, total RNA counts, and ribosomal read percentage) covariates were included. The model was trained for up to 400 epochs with early stopping and lr = 1e-3.

The scVI latent space was used as the basis for CD8^+^ T cell state discovery. A nearest-neighbor graph was constructed using the scVI embedding (n_neighbors = 15), followed by Leiden clustering (resolution = 0.5). This Leiden resolution was selected after reviewing clustering solutions across a local resolution sweep (resolution = 0.3–0.9) for cluster separation (silhouette score), marker coherence, donor representation, and overfragmentation. Resulting clusters were annotated manually using established CD8^+^ T cell marker expression and cluster-level differentially expressed genes. One mixed cluster was subclustered in the scVI latent space (n_neighbors = 15, resolution = 0.2) to resolve central memory CD8^+^ T cells from contaminating cytotoxic CD4^+^ T cells, and this cytotoxic CD4^+^ T cell subcluster was removed. Two additional clusters were excluded including one small single-sample cluster with discordant cell lineage marker expression and one rare nonspecific cluster comprising <0.1% of cells.

The final CD8^+^ T cell object contained 97,167 cells across seven annotated CD8^+^ T cell clusters. For marker visualization and quantification, expression values were normalized (target_sum = 1e4) and log-transformed.

### Proportionality analysis

Cell-state proportions were quantified at the donor level using Scanpro (v0.2) from final CD8^+^ T cell annotations. Where indicated, CD8^+^ T cell labels were transferred back to the parent PBMC object to assess proportions relative to total PBMCs.

### Differential gene expression, pathway analyses, and gene set enrichment

Cluster-enriched genes were identified using Scanpy (scanpy.tl.rank_genes_groups) with Wilcoxon rank-sum testing and Benjamini-Hochberg correction. Focused differential expression analyses between CD8^+^ T cell states were performed using donor-balanced iterative sampling to minimize bias from unequal donor contribution to variably abundant states. For each comparison, genes were retained only if detected in at least 5% of cells in either group to reduce sparse-expression noise. Up to 50 cells per donor per cell state were randomly sampled across 50 iterations, and Wilcoxon testing was performed for each iteration. Differential expression results were summarized using the median log2 fold change, test statistic, and adjusted P value.

Pathway activity was inferred using decoupler (v1.6.0) with univariate linear modeling and MSigDB Hallmark gene sets. Scores were calculated from normalized single-cell expression values and summarized at the donor and CD8^+^ T cell state level. Gene set enrichment analysis was performed using GSEApy (v1.1.4) with preranked statistics from the donor-balanced differential expression analysis and the KEGG 2021 Human gene set library.

### Metabolic flux balance analysis

Metabolic flux balance analysis was performed using Compass (v0.9.10.2). For each CD8^+^ T cell state, single-cell transcriptomes were normalized to counts per million. Compass was run with the Recon2 human metabolic reconstruction using the IBM CPLEX optimizer and penalty diffusion by k-nearest neighbors, lambda = 0, num_neighbors = 30, and microclusters of 25 cells.

Compass reaction penalty scores were negative log-transformed so that larger values represented greater inferred reaction flux potential. Low-confidence and invariant reactions were removed, and positive flux directions were retained for bidirectional reactions. Reaction-level flux potential was summarized by CD8^+^ T cell state using z-scores for visualization and Cohen’s d effect sizes for pairwise comparisons.

### Gene regulatory network analysis

Transcription factor activity was inferred using decoupler with univariate linear modeling and the human CollecTRI regulatory network. The network was filtered to target genes present in the CD8^+^ T cell object. Transcription factors were retained if they were detected in ≥3% of cells and had ≥15 target genes in the filtered network. Activity scores were calculated from normalized expression values and analyzed within CI effector CD8^+^ T cell states. For visualization, transcription factor activity was summarized as the median activity per donor and CD8^+^ T cell state, z-scored across states, and filtered to transcription factors with nonzero variance and significant cluster-level activity differences.

Focused regulatory comparisons between T_Eff-1_ and T_Eff-2_ were performed using donor-level median transcription factor activity among donors with at least 5 cells in both states. Differential transcription factor activity was tested using two-sided Mann-Whitney U tests with Benjamini-Hochberg correction. To estimate candidate gene target-level regulatory effects, significant transcription factors were mapped back to signed CollecTRI edges. Target genes were retained if detected in at least 5% of T_Eff-1_ or T_Eff-2_ cells. Each edge was weighted by the difference in transcription factor activity between T_Eff-2_ and T_Eff-1_ multiplied by the signed regulatory interaction, and gene target-level effects were summarized across convergent transcription factors. Network visualizations were generated with NetworkX (v3.5).

### Velocity and pseudotime analyses

Spliced and unspliced RNA count matrices were generated using the alevin-fry (v0.11.2) pipeline. A splici (spliced + intronic) reference was constructed from the GRCh38-2020-A genome FASTA and GTF using pyroe (v0.9.0) make-splici with a 151 bp read length and 5 bp flank trim, and indexed with salmon (v1.10.1). Reads from each of the 47 samples were mapped using salmon alevin in sketch mode with ISF library orientation and 10x Chromium barcode geometry. Cell barcodes were identified by knee calling, and quantification was performed with cr-like UMI resolution using a three-column transcript-to-gene map to enable USA (Unspliced/Spliced/Ambiguous) mode. Per-sample matrices were concatenated and transferred onto the annotated CD8^+^ T cell object by barcode matching.

RNA velocity was modeled using VeloVI (v0.3.1). Genes with fewer than 30 shared spliced and unspliced counts were removed, and neighborhood-smoothed spliced and unspliced counts were computed using the existing scVI neighbor graph to preserve the original embedding. The VeloVI preprocess_data function was applied to filter genes by steady-state model fit and scale smoothed counts to [0, 1], yielding 209 genes for model training. The model was initialized with default architecture (n_hidden = 256, n_latent = 10, n_layers = 1, dropout_rate = 0.1) and trained for up to 500 epochs with early stopping on validation ELBO. Velocity vectors and directional uncertainty were extracted from the trained model and projected onto the scVI UMAP embedding using scvelo (v0.3.4). Permutation scores were computed per gene across CD8^+^ T cell clusters to assess the consistency of spliced and unspliced count dynamics with transient kinetics.

Palantir pseudotime (v1.4.4) was computed from the scVI latent embedding with n_components = 5 and num_waypoints = 500, rooted in a T_N_ cell selected as the cell nearest to the T_N_ cluster centroid in scVI space. A CellRank2 (v2.0.7) PseudotimeKernel was constructed from Palantir pseudotime and combined with a ConnectivityKernel derived from the scVI neighbor graph at a ratio of 0.8:0.2. Random walks were simulated from T_N_ cells and projected onto the UMAP embedding (n_sims = 100, max_iter = 200, seed = 0). Macrostates were identified using the GPCCA estimator with Schur decomposition (n_components = 20) and 13 macrostates. Split macrostates corresponding to the same biological population were merged into unified terminal states prior to fate probability computation. Terminal states were defined as T_Eff-2_, T_Eff-1_, T_KIR_, and T_CMC1_ with T_N_ as the initial state.

To prevent clonal TCR variable gene expression from confounding lineage driver rankings, T cell receptor genes and pseudogenes were excluded using curated HGNC locus type annotations (locus_type: “T cell receptor gene” and “T cell receptor pseudogene”). Fate probability-weighted lineage driver analysis was performed using CellRank 2 compute_lineage_drivers on normalized expression values within T_EM_, T_Eff-1_, and T_Eff-2_ cells, comparing T_Eff-2_ and T_Eff-1_ lineages simultaneously. Driver genes were identified by Pearson correlation with fate probability and defined at q < 0.05 after Benjamini-Hochberg correction. For gene expression trend visualization, genes detected in fewer than 10 cells were removed and magic-impute (v3.0.0) was applied to the remaining gene set (name_list = “all_genes”, knn = 5, t = “auto”) to reduce dropout noise, yielding 8,092 genes. Generalized additive models were then fit to MAGIC-imputed expression weighted by lineage fate probability along Palantir pseudotime.

### Flow Cytometry

Cells were transferred to 96-well V-bottom plates (Corning, cat. no. 3894), pelleted at 300g for 5 min at 4°C, and washed twice in cold PBS. Viability and Fc receptor blocking were performed simultaneously in PBS for 30 min at 4°C using Ghost Dye Violet 450 Fixable Viability Dye (Cytek Biosciences, cat. no. 13-0863-T100, 1:20,000) and Human TruStain FcX (BioLegend, cat. no. 422302, 1:40). After washing in FACS buffer (PBS with 2% FBS), cells were incubated with surface antibodies (Supplemental Table 8) in FACS buffer for 30 min at 4°C and washed twice. Data were acquired on a Cytek Aurora (V-B-YG-R) and analyzed in FlowJo v.10.9.0 (Becton Dickinson & Company).

### CyTOF

CyTOF was performed as previously described (53, 54). Briefly, cryopreserved PBMCs were rapidly thawed in a 37°C water bath and recovered in RPMI 1640 supplemented with 10% FBS and DNAse I. Cells were washed and stained with rhodium-based Cell-ID viability reagent (Fluidigm) according to the manufacturer’s recommendations, followed by staining with metal-conjugated antibodies targeting cell-surface markers for 30 min at room temperature. Cells were then washed, fixed with 1.6% paraformaldehyde, permeabilized with ice-cold methanol, and stained with metal-conjugated antibodies targeting intracellular proteins. Cells were stained with iridium Cell-ID DNA intercalator (Fluidigm), washed with PBS and ultrapure water, resuspended with EQ Four Element Calibration Beads, and filtered before acquisition on a Helios CyTOF 3.0 mass cytometer at the Vanderbilt Cancer & Immunology Core. Data were collected as FCS files and normalized using bead-based normalization (55). Antibodies used for staining are listed in Supplemental Table 9.

### CyTOF Analysis

CyTOF data were analyzed in OMIQ software (Dotmatics). CD8^+^ T cells were identified by manual pre-gating on CD45^+^CD3^+^CD19^-^CD8^+^CD4^-^CD20^-^CD14^-^CD66b^-^Ki67^-^TCRγδ^-^ cells (Supplemental Fig. 4). Samples with ≥515 CD8^+^ T cells were subsampled to 515 cells; all cells from samples below this threshold (n=3) were included. UMAP dimensionality reduction was applied to the pre-gated subsampled data (Neighbors = 15, Minimum Distance = 0.4, Components = 2, Metric = Euclidean, Learning Rate = 1, Epochs = 200, Embedding Initialization = spectral, Random Seed = 7408). FlowSOM clustering was performed on all markers not used in the pre-gating (xdim = 12, ydim = 12, rlen = 10, Distance Metric = Euclidean, Consensus Metaclustering, k = 20, Random Seed = 9438). Expert annotation of metaclusters yielded a coarse 6-population and fine 11-population clustering, both derived from the same FlowSOM run by varying the degree of cluster merging during annotation. For the selected-marker re-clustering, UMAP and FlowSOM were repeated using the same parameters but restricted to the 10 markers we identified as most differentially expressed in CD8^+^ T cells from sample V-0055 (CD127, CD27, ICOS, CCR7, GLUT1, CD95, CD38, CytoC, CD57, PD-1). For the CCR7^-^selected-marker analysis, the pre-gate was extended to additionally exclude CCR7^+^ cells, samples were re-subsampled to 374 cells (files with <374 cells (n=4) had all cells included), and the previous pipeline was repeated.

Per-sample CyTOF cluster proportions were exported from OMIQ and analyzed in R (v4.4.1). Cluster proportions were compared between HC and CI using Mann-Whitney U test (wilcox.test, base R). Pearson correlation (cor.test, base R) and Lin’s concordance correlation coefficient (DescTools v0.99.60) between paired per-sample CyTOF cluster proportions and scRNA-seq CD8^+^ T cell state proportions were computed on raw and log_10_-transformed proportions, with significant associations defined at p<0.05. Results were visualized using ggplot2 (v4.0.1) and ComplexHeatmap (v2.20.0). The CIRTUS algorithm was used for CyTOF cell state discovery. Pre-gated, subsampled CD8^+^ T cells were exported from OMIQ as FCS files. CITRUS (v0.8) was then run in R (v4.4.1) with family=“continuous”, modelTypes=“sam”, nFolds=1, fileSampleSize=515, minimumClusterSizePercent=0.5, transformCofactor=5, and featureType=“abundances” on all panel markers not used in the pre-gating. Both raw and logit-transformed T_Eff-2_ proportions were evaluated as endpoints against scRNA-seq-derived per-sample T_Eff-2_ proportions; for the logit transformation, exact zeros were replaced with 0.001 prior to transformation.

### External bronchoalveolar lavage CD8^+^ T cell analysis

A public single-cell RNA sequencing dataset of CD3-enriched bronchoalveolar lavage cells (GSE167118) was analyzed as an external tissue cohort. Cells with <400 or >5,000 detected RNA features or >20% mitochondrial reads were excluded. Doublets were identified and removed using DoubletDetection (v4.2). CD8-enriched T cells were identified from the CD3-enriched object by coarse clustering and *CD8A*/*CD8B* expression, then reanalyzed using scVI with 2,000 highly variable genes. A nearest-neighbor graph was constructed from the scVI latent space (n_neighbors = 20) followed by Leiden clustering (resolution = 0.4). One small low-quality cluster with low feature counts, high mitochondrial content, and mitochondrial gene-enriched marker expression was removed as putatively dead/dying cells before final annotation.

BAL CD8^+^ T cell states were annotated manually using canonical marker expression and cluster-level differentially expressed genes. Cell-state proportions were quantified at the donor level using Scanpro. A T_Eff-2_ gene module was derived from the primary PBMC CD8^+^ T cell dataset by identifying genes enriched in T_Eff-2_ compared with all other CD8^+^ T cell states (adjusted p < 0.05, log2 fold change > 0.5, expression in ≥20% of T_Eff-2_ cells) and absent from the corresponding marker lists of other CD8^+^ T cell states (Supplemental Table 7). This module was scored in the BAL CD8^+^ T cell clusters using Scanpy (score_genes). Hallmark pathway activity was inferred using decoupler with univariate linear modeling, as described above.

### External PBMC validation analysis

A public single-cell RNA sequencing dataset of PBMCs from adult patients with sepsis and healthy controls (GSE279452) was analyzed as an external validation cohort. Two samples (Abd-S8 and Abd-S80; GEO accessions GSM8571120 and GSM8571121) with ambiguous metadata-to-matrix pairing at the time of download were excluded, leaving 7 healthy control and 91 adult sepsis samples in the analysis. Cells with <400 or >5,000 detected RNA features or >20% mitochondrial reads were excluded, genes detected in fewer than 3 cells were removed, and doublets were identified and removed using DoubletDetection.

Major PBMC lineages were first identified using scVI with 3,000 highly variable genes. CD8^+^ T cells were identified by coarse clustering and *CD8A*/*CD8B* expression, then integrated with the annotated discovery CD8^+^ T cell dataset using scANVI over shared highly variable genes. The scANVI model was specified with n_hidden = 128, n_latent = 10, n_layers = 1, and gene_likelihood = “zinb” and trained for 100 epochs. This produced a validation CD8^+^ T cell latent space in which scANVI assigned validation cohort cells to discovery-defined states with prediction probabilities. Downstream analyses were then performed within the external validation cohort.

Cell-state proportions, T_Eff-2_ versus T_Eff-1_ differential expression, and KEGG gene set enrichment were performed as described above. Mortality associations were tested using logistic regression (statsmodels v0.14.1), comparing the highest T_Eff-2_ quintile to the lower four quintiles in unadjusted and age/sex-adjusted models for 14-day, 28-day, and 90-day mortality.

### Statistical analyses

Statistical analyses were performed in Python using scipy.stats (v1.11.4) and statsmodels (v0.14.1). Two-group comparisons used two-sided Mann-Whitney U tests, continuous associations used Spearman rank correlation, and ordered PaO_2_/FiO_2_ categories were tested using the Jonckheere-Terpstra trend test. LOWESS curves with 1,000 bootstrap resamples were used for visualization where indicated. P values were adjusted using the Benjamini-Hochberg false discovery rate method where indicated.

Cell-state proportions were analyzed at the donor level using Scanpro empirical Bayes moderated testing of logit- or arcsin-transformed proportions. Differential expression used Wilcoxon rank-sum testing with Benjamini-Hochberg correction. Gene set enrichment used GSEApy preranked analysis with a permutation-based weighted Kolmogorov-Smirnov enrichment statistic and false discovery rate correction.

External validation mortality analyses used binary logistic regression comparing the highest T_Eff-2_ abundance quintile with the lower four quintiles in unadjusted and age/sex-adjusted models. Unless otherwise indicated, each data point represents an individual donor. No statistical method was used to predetermine sample size; analyses used all eligible samples available from the discovery and validation datasets after quality control and predefined exclusions. Clinical variables unavailable in the electronic health record were omitted from analyses requiring those values. Investigators were not blinded to sample group assignments during analysis.

### Study Approval

All studies were reviewed and approved by the Vanderbilt Institutional Review Board under protocols 211462, 191562, and 250934. Written informed consent was obtained from all participants or legally authorized surrogates.

### Data and code availability

The full processed PBMC single-cell RNA-sequencing dataset from which the discovery CD8^+^ T cell data were derived was generated previously and is available through the Gene Expression Omnibus under accession GSE290679. Raw sequencing data and patient-level metadata are available through dbGaP under controlled access, with ICU participant data under phs004258.v1.p1 and healthy control data under phs004377.v1.p1, in accordance with participant consent and patient privacy protections. External datasets were obtained from GEO under accessions GSE167118 (CD3-enriched bronchoalveolar lavage single-cell RNA-sequencing dataset) and GSE279452 (adult sepsis and healthy control PBMC single-cell RNA-sequencing dataset).

Processed CD8^+^ T cell analysis objects generated for this study, including the discovery PBMC CD8^+^ T cell object, external bronchoalveolar lavage CD8^+^ T cell object, and external PBMC validation CD8^+^ T cell object, will be made available through GEO. Analysis code and notebooks will be deposited at GitHub upon publication.

## Supporting information

Table1_PatientDemographics

## Author Contributions

C.M.N., L.B.W., and M.T.S. designed the research and prepared the manuscript with contributions from all other authors. C.M.N., M.T.S., S.N.O., and V.E.K. performed the research and analyzed data. D.C.N. and J.C.R. provided essential intellectual support. J.A.B. and L.B.W. provided essential expertise and clinical samples. C.E.R., J.M.I, and J.Y.C. provided key technical support. E.L.M., C.Y.S., and M.T.S. collected and/or processed clinical samples.

## Acknowledgements

We sincerely thank the patients for their participation and for generously providing the biospecimens essential to this study. We appreciate the clinical staff who collaborated with our research team and made this work possible. We are grateful for insightful discussions with members of the laboratories of Dr. Lorraine Ware, Dr. Julie Bastarache, Dr. Ciarra Shaver, Dr. Jeffrey Rathmell, and Dr. Kimryn Rathmell. We also appreciate the support of Dr. Angela Jones and staff at the Vanderbilt Technologies for Advanced Genomics (VANTAGE) core.

