## Supplemental_Figures for "Critical illness expands a transcriptionally distinct hypometabolic CD8^+^ T effector program associated with respiratory failure and mortality"

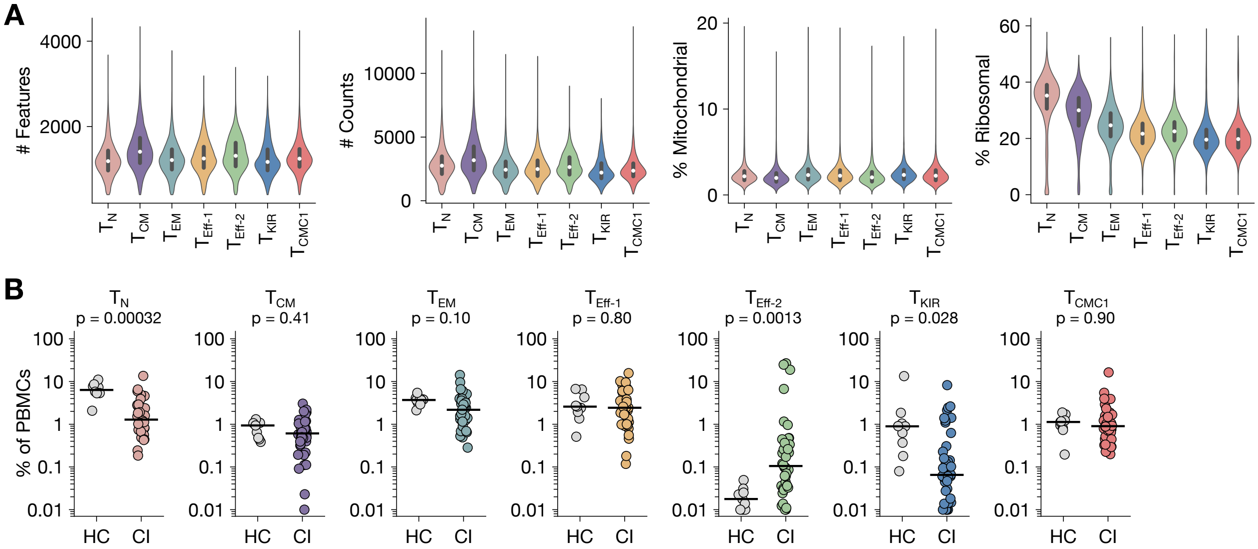


**Supplemental Figure 1: scRNA-seq $CD8^+$ T cell cluster quality control metrics and $CD8^+$ T cell subset frequencies among all PBMCs. (A)** Number of detected unique genes (features), total counts, percentage of mitochondrial reads, and percentage of ribosomal reads per cell after preprocessing. **(B)** Per participant frequency of each $CD8^+$ T cell cluster among all PBMCs in HC and CI patients. Statistical analysis was performed using the Empirical Bayes moderated T-test **(B)**. Each datapoint represents an individual research participant.

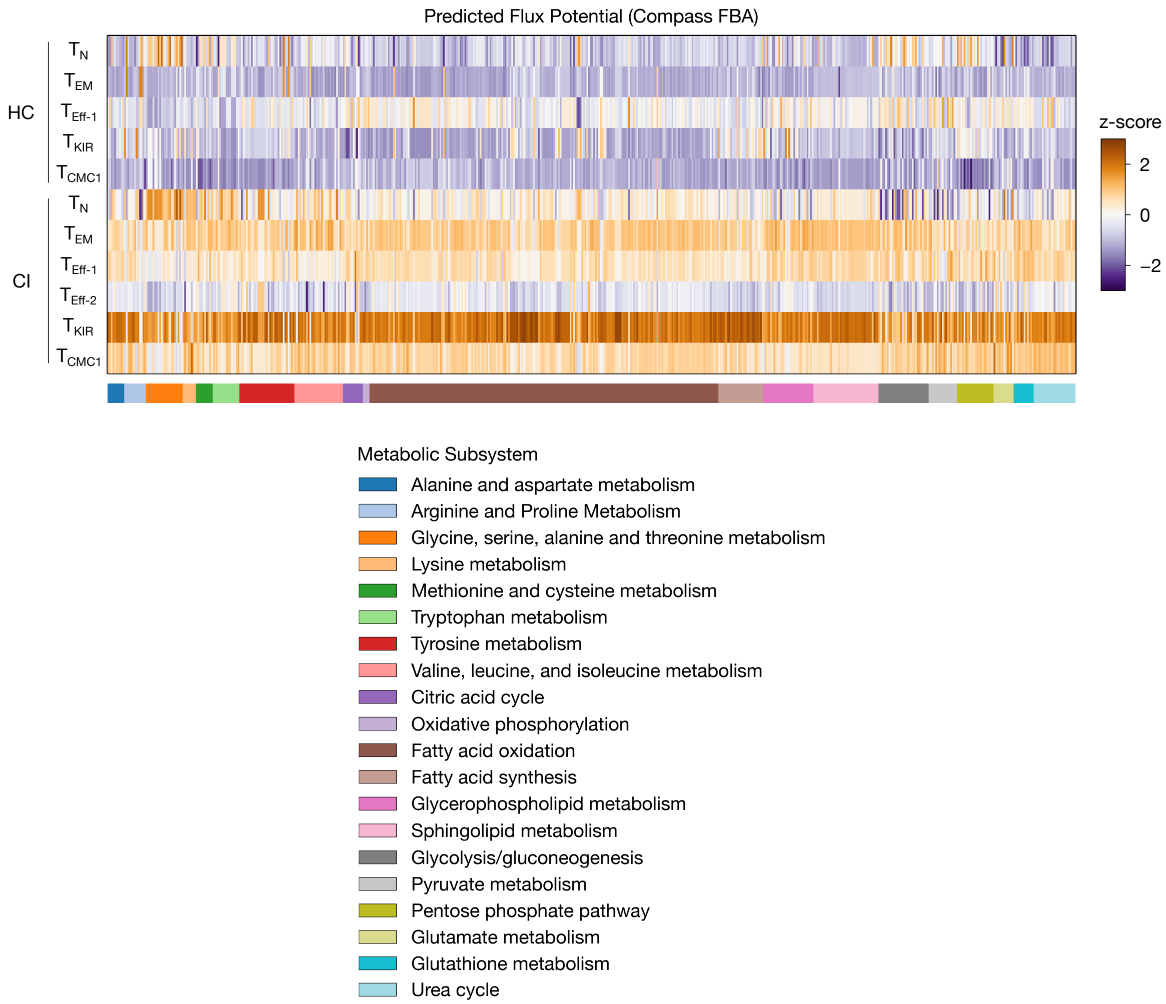


**Supplemental Figure 2: Compass flux balance analysis of metabolic reaction flux across all CD8$^+$ T cell subsets.** Flux balance analysis (FBA) via Compass as a heatmap showing *in silico* flux potential for individual metabolic reactions scaled by standard z-scoring from HC and CI CD8$^+$ $T_N$, $T_{EM}$, $T_{Eff-1}$, $T_{Eff-2}$, $T_{KIR}$, and $T_{CMC1}$ cells, where sufficient cells for analysis were present. Each column represents an individual metabolic reaction in the Recon2 network. Larger values indicate higher predicted flux.

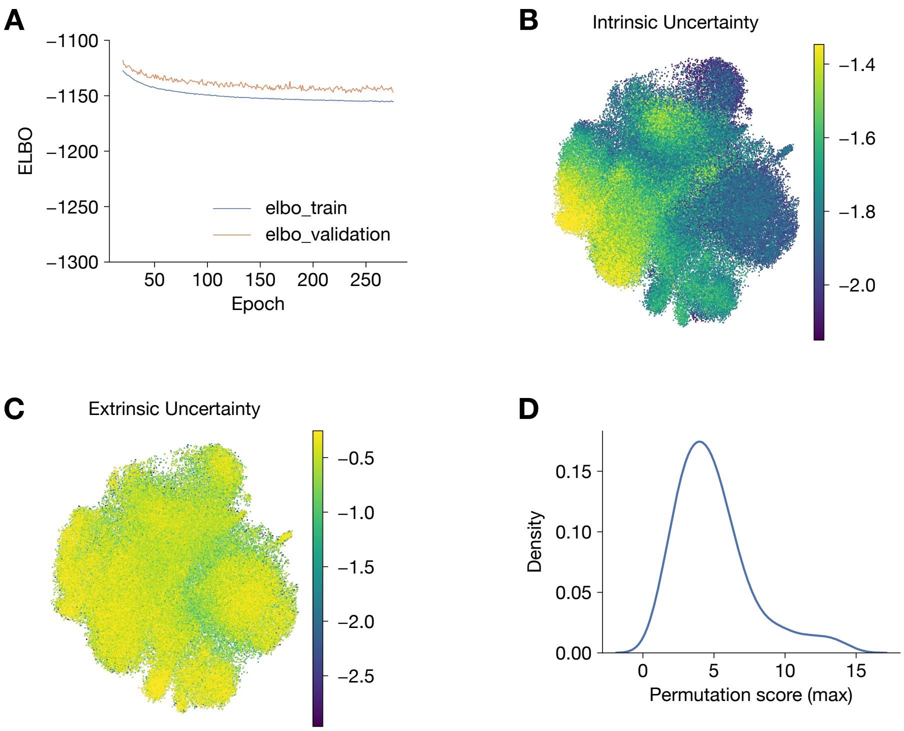


**Supplemental Figure 3: VeloVI model quality control metrics. (A)** Training and validation evidence lower bound (ELBO) curves across epochs during VeloVI model training. **(B)** UMAP visualization of intrinsic directional uncertainty, shown as $\log_{10}$-transformed directional cosine similarity variance across cells. **(C)** UMAP visualization of extrinsic directional uncertainty, shown as $\log_{10}$-transformed directional cosine similarity variance across cells. **(D)** Permutation scores across genes, quantifying the increase in VeloVI model fit error upon permutation of spliced and unspliced abundances. Higher scores indicate genes whose dynamics are consistent with transient rather than steady-state behavior.

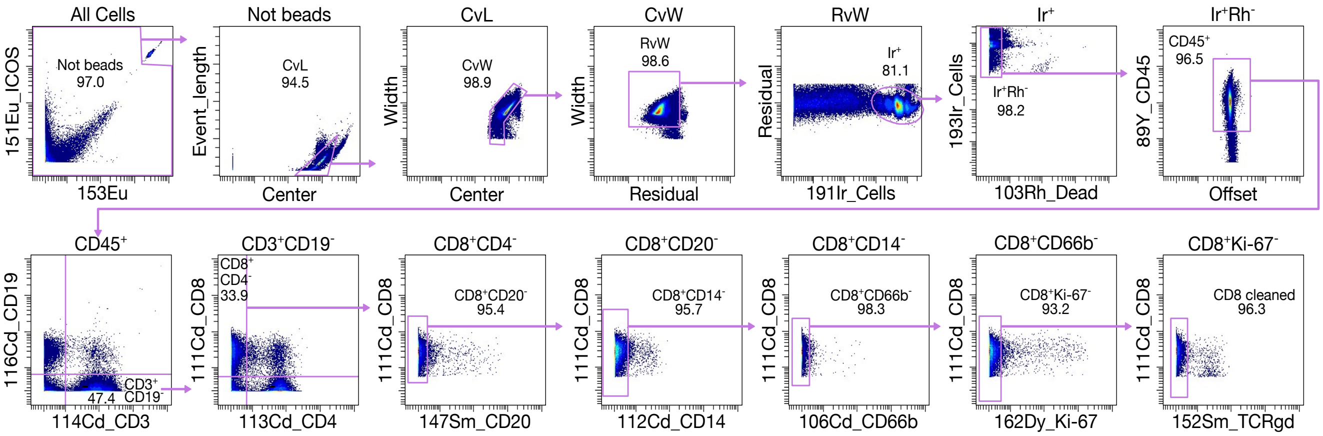


**Supplemental Figure 4: Representative CyTOF gating strategy.** Gating strategy for the identification of cleaned CD8$^{+}$ T cells (CD45$^{+}$CD3$^{+}$CD19$^{-}$CD8$^{+}$CD4$^{-}$CD20$^{-}$CD14$^{-}$CD66b$^{-}$Ki67$^{-}$TCRγδ$^{-}$) by CyTOF for downstream clustering.

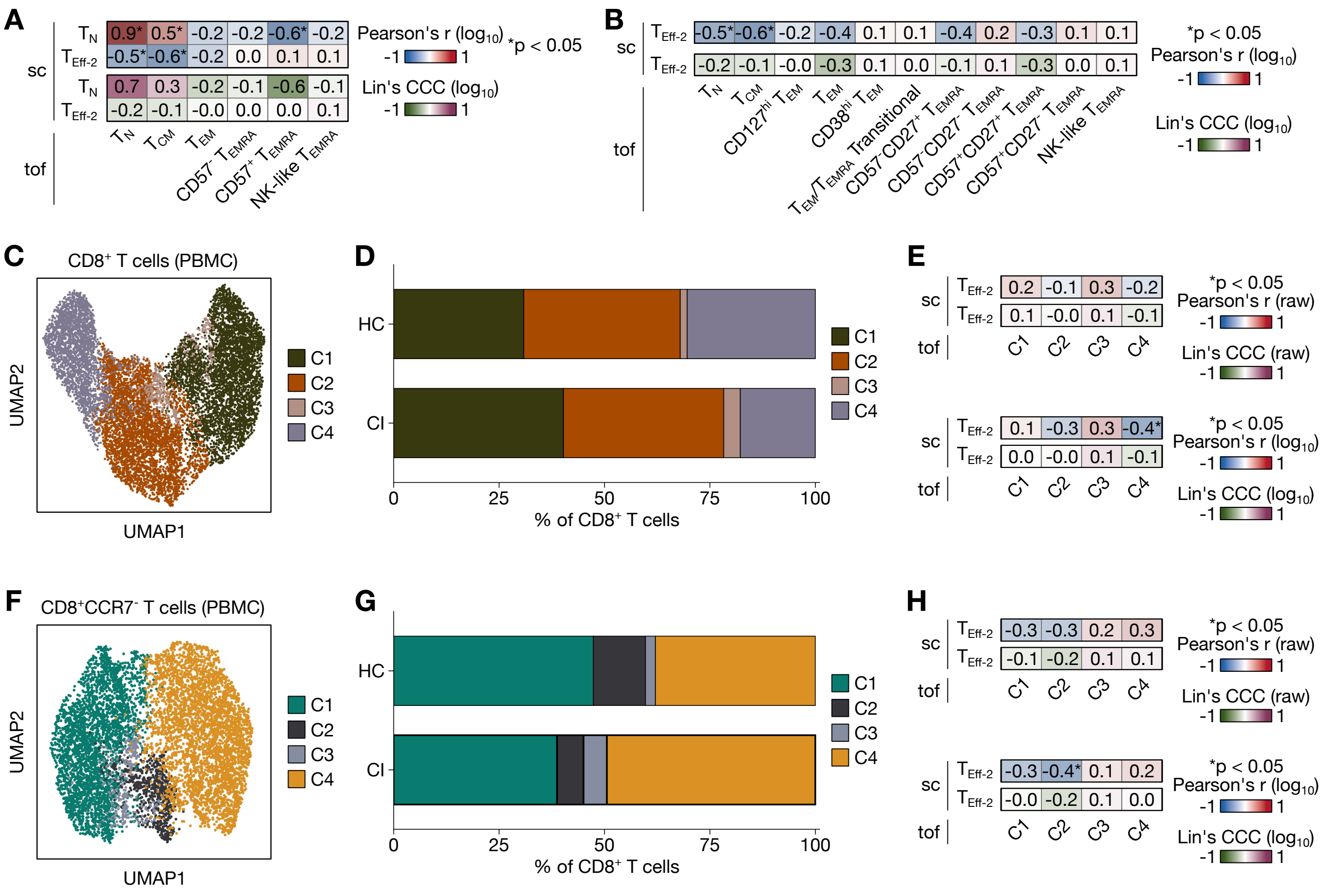

**Supplemental Figure 5: Additional protein-level analyses fail to resolve CD8$^+$ T$_{Eff-2}$. (A)** Pearson correlation coefficients and Lin's CCC of scRNA-seq T$_N$ and T$_{Eff-2}$ cluster proportions compared to all CyTOF cluster proportions from Fig. 6J, shown for log$_{10}$-transformed data. **(B)** Pearson correlation coefficients and Lin's CCC of scRNA-seq T$_{Eff-2}$ cluster proportions compared to all high-resolution CyTOF cluster proportions from Fig. 6N, shown for log$_{10}$-transformed data. **(C)** UMAP visualization of CyTOF CD8$^+$ T cells clustered on the 10 markers most distinguishing all CD8$^+$ T cells in the sample with the highest scRNA-seq T$_{Eff-2}$ frequency in the CyTOF dataset (V-0055). **(D)** Mean proportion of each cluster among all CD8$^+$ T cells in HC and CI patients. **(E)** Heatmap of Pearson correlation coefficients and Lin's CCC of scRNA-seq T$_{Eff-2}$ cluster proportions compared to all CyTOF cluster proportions from **(C)**, shown for raw and log$_{10}$-transformed data. **(F)** UMAP visualization of CyTOF CD8$^+$CCR7$^-$ T cells clustered on the 9 remaining markers (excluding CCR7) most distinguishing the sample with the highest scRNA-seq T$_{Eff-2}$ frequency (V-0055). **(G)** Mean proportion of each cluster among all CD8$^+$CCR7$^-$ T cells in HC and CI patients. **(H)** Heatmap of Pearson correlation coefficients and Lin's CCC of scRNA-seq T$_{Eff-2}$ cluster proportions compared to all CyTOF cluster proportions from **(F)**, shown for raw and log$_{10}$-transformed data. Statistical testing was performed using the Pearson correlation coefficient with two-sided p value with significant p values < 0.05 denoted **(A, B, E, H)**. Lin's concordance correlation coefficient were also reported **(A, B, E, H)**. sc = scRNA-seq; tof = CyTOF.

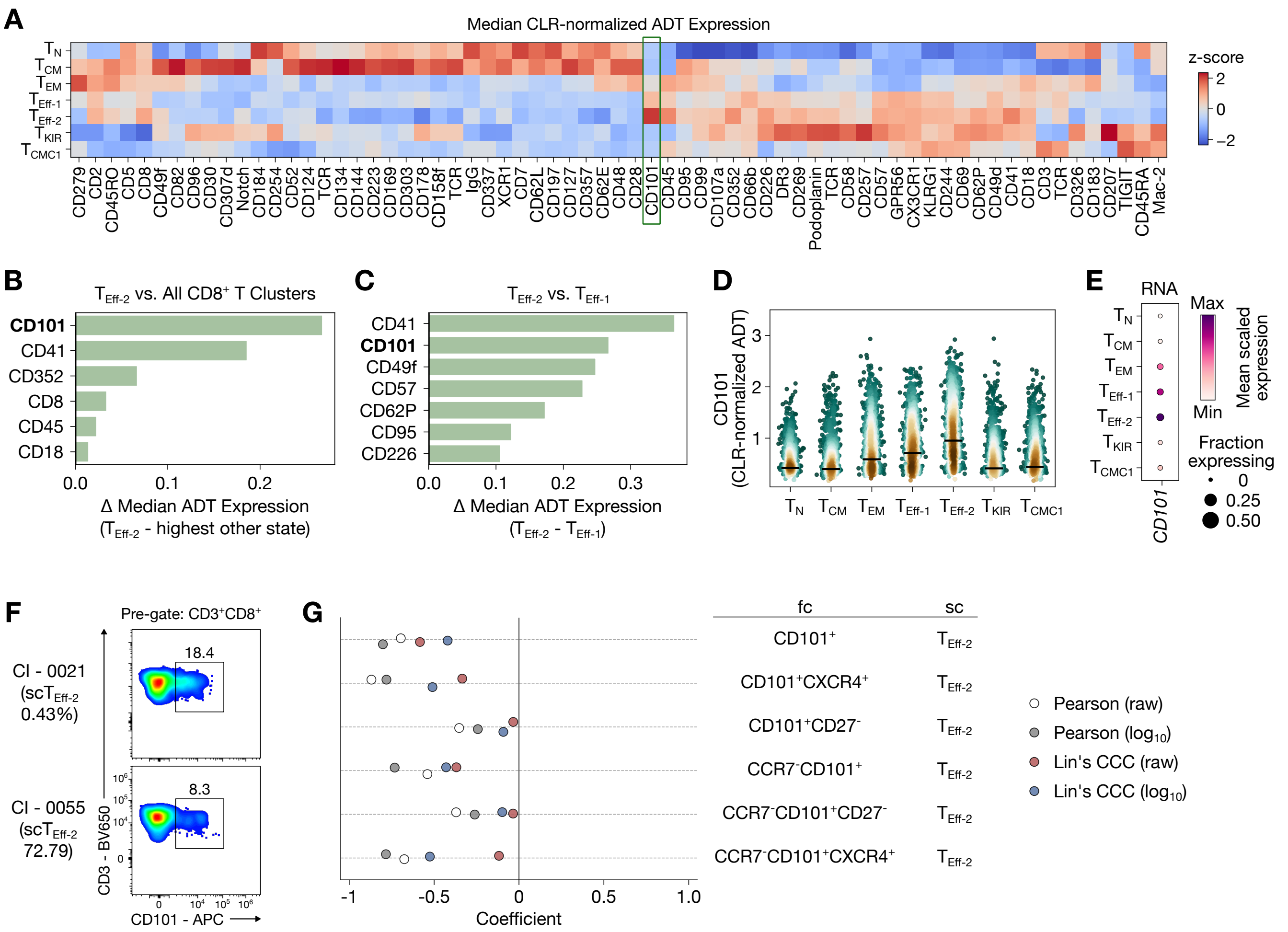


**Supplemental Figure 6: CITE-seq analysis of $T_{Eff\text{-}2}$.** **(A)** Heatmap showing z-scored, center log ratio (CLR) normalized antibody-derived tag (ADT) median abundance across $CD8^+$ T cell clusters. Markers shown represent those with median ADT signal above isotype control in ≥50% of cells in at least one $CD8^+$ T cell cluster. **(B)** Effect size of median ADT abundance for $T_{Eff\text{-}2}$ versus all other $CD8^+$ T cell clusters and **(C)** $T_{Eff\text{-}2}$ versus $T_{Eff\text{-}1}$. **(D)** CD101 ADT expression (CLR-normalized) per cell for each $CD8^+$ T cell cluster. **(E)** CD101 per-cell transcript abundance for each $CD8^+$ T cell cluster. **(F)** Representative flow plots of $CD101^+CD8^+$ T cells. **(G)** Pooled frequencies and correlations (Pearson's and Lin's CCC) of $CD101^+CD8^+$ T cells, including sensitivity analyses performed with additional gating for CXCR4, CD27, and/or CCR7 as specified.

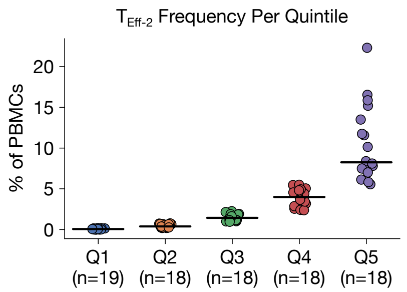


**Supplemental Figure 7: $T_{Eff\text{-}2}$ frequency per quintile.** Percent $T_{Eff\text{-}2}$ of all PBMCs stratified by quintile.
